# Targeted genome mining with GATOR-GC maps the evolutionary landscape of biosynthetic diversity

**DOI:** 10.1101/2025.02.24.639861

**Authors:** José D. D. Cediel-Becerra, Andrés Cumsille, Sebastian Guerra, Yousong Ding, Valérie de Crécy-Lagard, Marc G. Chevrette

## Abstract

Gene clusters, groups of physically adjacent genes that work collectively, are pivotal to bacterial fitness and valuable in biotechnology and medicine. While various genome mining tools can identify and characterize gene clusters, they often overlook their evolutionary diversity, a crucial factor in revealing novel cluster functions and applications. To address this gap, we developed GATOR-GC, a targeted genome mining tool that enables comprehensive and flexible exploration of gene clusters in a single execution. We show that GATOR-GC identified a diversity of over 4 million gene clusters similar to experimentally validated biosynthetic gene clusters (BGCs) that other tools fail to detect. To highlight the utility of GATOR-GC, we identified previously uncharacterized co-occurring conserved genes potentially involved in mycosporine-like amino acid biosynthesis and mapped the taxonomic and evolutionary patterns of genomic islands that modify DNA with 7-deazapurines. Additionally, with its proximity-weighted similarity scoring, GATOR-GC successfully differentiated BGCs of the FK-family of metabolites (e.g., rapamycin, FK506/520) according to their chemistries. We anticipate GATOR-GC will be a valuable tool to assess gene cluster diversity for targeted, exploratory, and flexible genome mining. GATOR-GC is available at https://github.com/chevrettelab/gator-gc.

**Graphical abstract:** 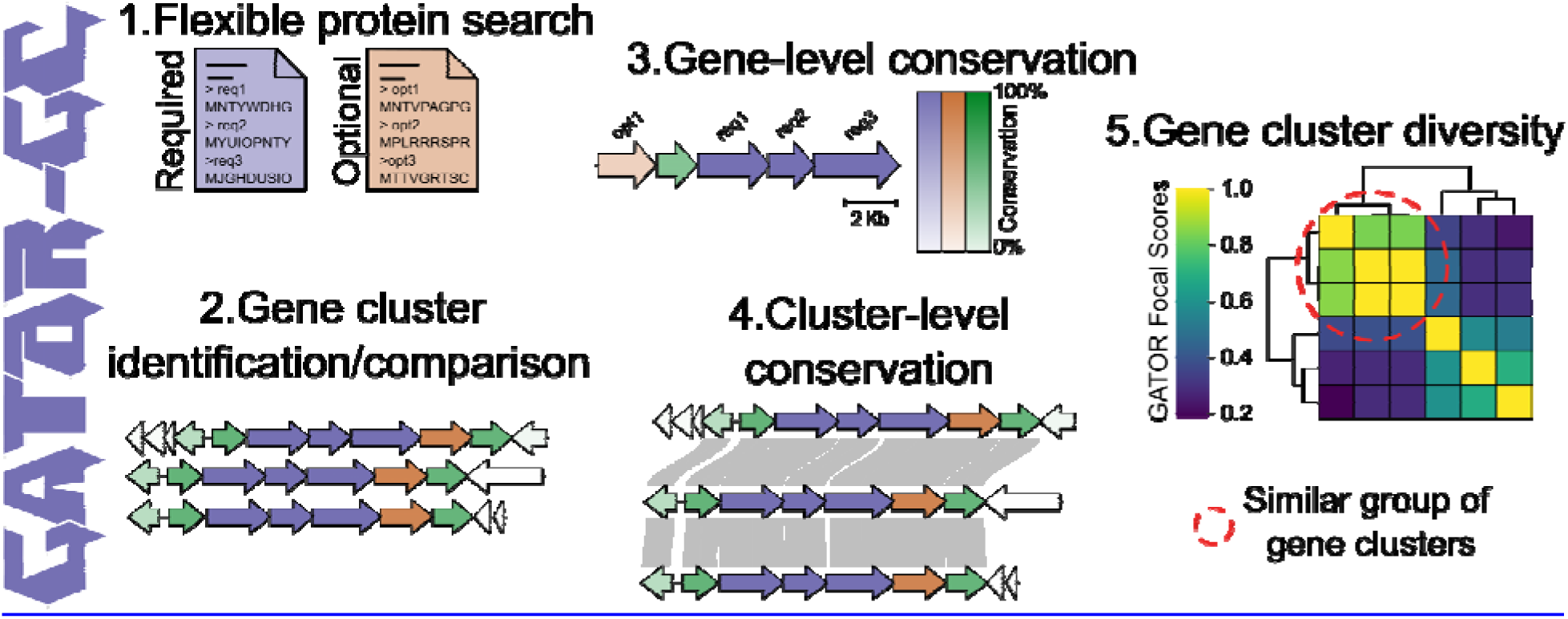

## Introduction

Genes that function collectively are often physically adjacent in bacterial genomes, forming gene clusters whose products play diverse and important roles (1, 2), including in environmental sensing (3), pathogenicity (4), nutrient acquisition (5), energy conversion (6), and the biosynthesis of bioactive molecules (7). For instance, the type III secretion system in *Salmonella typhimurium* is encoded by a gene cluster of at least 25 genes, which assemble its structural components and produce effector proteins that are injected into host cells to promote pathogenesis (8). Biosynthetic gene clusters (BGCs) direct the assembly of secondary metabolites that mediate interspecies interactions (9). BGCs are of particular societal interest because their products have been used as antimicrobials, biopesticides, photoprotectants, and crop-protectant agents, among others (10–13). Exploring the diversity of gene clusters is crucial not only for understanding their roles in bacterial fitness but also for uncovering potential applications in biotechnology and medicine.

Various genome mining tools have been developed to identify and analyze gene clusters, using approaches including machine learning (7, 14–18), comparative genomics (19–23), evolutionary frameworks (24, 25), hybrid workflows (26), and rule-based methods (27, 28). For instance, cblaster (23) enables the detection of gene clusters by querying protein sequences or PFAM profile identifiers, while CORASON (29) and clinker (30) support downstream comparative analysis and gene cluster visualization. The widely used BGC mining tool antiSMASH (27) employs pre-defined Hidden Markov Models (HMM) profiles of well-characterized biosynthetic genes and domains (e.g., non-ribosomal peptide synthetases [NRPSs] and polyketide synthases [PKSs]), to classify BGCs based on pathway-specific rules. Complementary tools like BiG-SCAPE (29) and BiG-SLICE (31) can then group BGCs with similar genomic architectures into gene cluster families. These and other genome mining tools have advanced the cataloging of bacterial BGCs in different databases leading to the computational prediction of tens of millions of BGCs (32–36), of which thousands have been experimentally validated and archived in the Minimum Information about a Biosynthetic Gene cluster (MIBiG, (36, 37)) data repository.

Despite this progress, current genome mining tools have limitations in exploring gene cluster diversity. While antiSMASH excels at identifying BGCs within an individual genome of interest, it was not designed to find or compare similar BGCs across large sets of many genomes. Most tools (e.g., cblaster and CAGECAT) lack all-vs-all gene cluster comparison capabilities, a critical feature for comprehensive comparative analysis. While these tools generate gene cluster diagrams with annotations, they do not visualize gene conservation, which is essential for identifying evolutionary patterns and refining query protein searches for more speculative hypotheses. For example, if a particular gene is consistently present across gene clusters from the same taxonomic group, its conservation may suggest functional importance and warrant further investigation. Currently, generating such hypotheses remains a manual process that requires chaining different tools together and/or the development of custom bioinformatic analyses.

Most workflows identify gene clusters based on sets of genes or domains that must all be present, lacking the flexibility to mark certain queries as optional (i.e., not strictly required). Incorporating the ability to designate queries as either required or optional would make gene cluster searches more flexible for exploratory research and conducive to iterative refinement. Although ClusterTools (38) partially addresses this issue, its scope is confined to BGCs (rather than all gene clusters), and its source code has not been updated in many years.

In addition, automated deduplication of gene clusters via genomic synteny evaluation is absent in existing tools, preventing the efficient elimination of redundant calculations during all-vs-all comparisons. Lastly, most workflows require chaining together multiple tools, and no single tool currently integrates flexible gene cluster searches, all-vs-all cluster comparisons, automated deduplication, annotation, and comprehensive visualizations in a single execution.

Here, we introduce GATOR-GC (Genomic Assessment Tool for Orthologous Regions and Gene Clusters) to address these gaps. GATOR-GC is a targeted genome mining tool designed to explore gene cluster diversity and conservation in a single execution. GATOR-GC allows users to classify proteins as either required or optional, providing flexibility for strategic and exploratory searches. It computes all-vs-all gene cluster comparisons and introduces GATOR Focal Scores (GFS), a metric that quantifies the similarity between gene clusters based on gene proximity to required and optional proteins, providing an evolutionary rationale for cluster boundary predictions and facilitating automated deduplication of identical clusters. Moreover, GATOR-GC outputs annotated GenBank files and generates informative gene cluster visualizations, including (i) a heatmap summarizing gene cluster diversity, (ii) gene conservation cluster diagrams to visualize gene-level conservation, and (iii) neighborhood diagrams to visualize cluster-level architectural conservation (or lack thereof). Available at https://github.com/chevrettelab/gator-gc, GATOR-GC is easily accessible and can be installed via conda.

To showcase the utility of GATOR-GC, we conducted a global analysis of all bacterial MIBiG v3.0 BGCs using antiSMASH v7.0. Within MIBiG, a total of 279 BGCs were unable to be detected and characterized by antiSMASH. To characterize the evolutionary landscape of this unexplored biosynthetic diversity, we used GATOR-GC to search ∼1.92M genomes and identified over 4 million gene clusters similar to those antiSMASH-missed BGCs. Further, we demonstrate GATOR-GC’s versatility across different gene clusters, including those encoding for known secondary metabolites and genomic islands, highlighting the utility of GATOR gene conservation diagrams, the optional protein search feature for describing gene cluster architectures, and the GFS system of mapping gene cluster diversity. Our findings demonstrate GATOR-GC’s effectiveness in exploring gene cluster diversity and variability, uncovering novel patterns of enzyme conservation, and investigating evolutionary dynamics in an exploratory, flexible, and iterative manner.

## Materials and Methods

### PRE-GATOR-GC Algorithm Implementation

#### Diamond Database

User-provided input genomes in GenBank format were parsed via SeqIO (BioPython v1.81) to generate a protein database. This process involved extracting coding sequences (CDSs) and their feature qualifiers, such as gene coordinates, strand orientation, locus tags, contig ID, and protein annotations. In cases where the translation feature qualifier was absent, amino acid sequences were translated from the DNA sequences. The headers of the resulting protein FASTA database were structured to include the genomic context qualifiers. This protein database was compiled into a DIAMOND database with DIAMOND v2.1.6 (39).

#### Modular Domains Database

Modular protein domains from nonribosomal peptide synthetases (NRPS; Adenylation [A] and Condensation [C] domains) and polyketide synthases (PKS; Acyltransferase [AT], and Ketosynthase [KS] domains) were identified by analyzing the protein database using HMM profiles from antiSMASH v7.0 (27) with the hmmsearch algorithm from HMMER v3.3.2 (40) at a default e-value threshold of 1e-4.

### GATOR-GC Algorithm Implementation

#### Categorization of Modular Protein Domains

A rule-based approach was used to categorize proteins in NRPS, PKS, and hybrids (NRPS-PKS). Briefly, proteins with at least one A domain and at least one C domain were categorized as NRPS, whereas those containing at least one AT domain and at least one KS domain were classified as PKS. Proteins classified as both NRPS and PKS were labeled as NRPS-PKS hybrids.

#### Screening for Modular Domains

Queries in protein FASTA format, designated either required or optional, were screened against HMM profiles of modular domains in NRPS and PKS using hmmsearch parameters as previously defined. Queries categorized as NRPS, PKS, or hybrids were separated from non-modular queries. In these cases, NRPS, PKS, or hybrids were instead searched by hmmsearch as previously described (see Modular Domains Database, above).

#### Protein Dictionary

Non-modular queries were aligned against the DIAMOND database generated by PRE-GATOR-GC, using DIAMOND v2.1.6 with default parameters for query cover (70%) and protein percentage identity (35%). These parameters are adjustable via command line arguments to accommodate user-specific requirements. Hits for modular queries (e.g., NRPS, PKS, hybrids) were retrieved from the modular domains database generated by PRE-GATOR-GC. Hits for both modular and non-modular protein queries were stored in a Python dictionary for subsequent identification of gene clusters.

#### Estimation of Typical BGC Intergenic Distances

Coding sequences (CDSs) from the bacterial MIBiG v3.0 (37) dataset were parsed to extract relevant feature qualifiers such as gene coordinates, locus tags, and protein annotations. For each MIBiG BGC, all intergenic distances between genes were computed based on the genomic space between the start and end positions. The default value for the “required_distance” command-line argument in GATOR-GC was set using the 95th percentile of these genomic distances (85.9 Kb, Supplementary Figure S1), reflecting distances observed in experimentally validated BGCs.

#### Gene Cluster Identification

Three conditions must be met to identify and define gene clusters from the protein dictionary: i) all user-required query proteins must be present on the same contig, ii) the distance between user-required query proteins must comply with the default distance cutoff value of 85.9 Kb (as defined above), and iii) a default distance extension, termed “window extension”, of 10 Kb upstream and downstream of the gene cluster boundaries was applied. Both required distance and window extension threshold values are adjustable via command line arguments to accommodate user-specific requirements. These gene clusters are collectively referred to henceforth as “GATOR windows”.

#### Annotation of GATOR Windows

GenBank files were generated for each identified GATOR window. Genome feature qualifiers were added for CDSs that had hits in the PRE-GATOR-GC DIAMOND Database. This included user-defined query protein type, designated as “gator_query” (i.e., required, optional, or None), the DIAMOND database hit labeled as “gator_hit”, and a Boolean label indicating NRPS or PKS, “gator_nrps” and “gator_pks”, respectively. A Boolean feature qualifier named “contig_edge” was created for all CDSs to mark genes located within 500bp of a contig edge. A table of all output was generated, containing the hits for the DIAMOND database, feature qualifiers, window identifiers, and genomic context information of the hits (e.g., gene coordinates, loci names, and contig). The hits were organized by the increasing order of window identifiers, with their genes arranged by their start positions. GenBank files were created under the folder “windows_genbanks” and were parsed to generate the protein and DIAMOND databases using the same module of PRE-GATOR-GC.

#### GATOR Window Deduplication

To avoid redundant calculations in the subsequent steps, identical GATOR windows to the GATOR Focal Window (GFW) go through a deduplication step. A GFW is defined as a window that is used to perform similarity comparisons to all other GATOR windows. Initially, GFWs were sorted by genomic length in descending order to determine the sequence for the deduplication process. GFW with the largest genomic length was first used to compute the GATOR Focal Scores (see Gene-level presence-absence and GATOR Focal Scores (GFS) sections below). GATOR windows with a GFS value of 1.0 were deemed identical and removed from the protein database, and their identifiers were stored in the deduplicated dictionary to track the GFW and their identical counterparts. The next largest GFW was used to continue with the deduplication process using the updated protein database. This procedure was repeated iteratively until all GATOR windows were processed and deduplicated. The deduplication results were saved in the “deduplication_process” folder. This final set of deduplication GATOR windows was used to create the GATOR-GC figures (See below).

#### Gene-level Presence-absence

A FASTA protein file from the GFW was used to query the protein database, either complete or the deduplicated subset via DIAMOND v2.1.6 with the same parameters described above (see Protein Dictionary, above). The output table from DIAMOND was then parsed out to generate presence-absence tables at the gene level for the GFW. These tables were exported in CSV format in the directory “presence_absence”.

#### GATOR Focal Score Calculations

To compare one GATOR window to another, the presence or absence of all genes in the GFW is evaluated against all other windows. Weights are applied to genes based on their proximity to the user-defined query proteins (i.e., required, and optional) within the GFW, using a decaying normal distribution. The maximum amplitude of this distribution (value of 1) is assigned to the positions of these user query proteins, with genes closer to them receiving higher weights and those farther away receiving lower weights. If a gene receives multiple weights from different required or optional genes, the highest weight is used. The resulting values were summed across genes in each window. To calculate the GFS values, each window’s summation value was divided by the summation value of the GFW. These GFSs ranged from 0.0 to 1.0, with GFS values close to 1.0 suggesting high similarity to the GFW, while values closer to 0.0 indicating low similarity. The resulting tables of these GFS values were exported in CSV format under the directory named “gator_scores”.

#### Clustered Heatmap

GFS values of all-vs-all windows were concatenated to construct a matrix, where each row represented a GATOR window, while each column represented a GFW. Pairwise Euclidean distances were computed using the SciPy v1.10.1 library and hierarchical clustering was performed on the distance matrix using complete linkage as the clustering method. The resulting dendrogram was plotted using Matplotlib v3.8.0 library, and a heatmap of the clustered GFS matrix was created employing the seaborn v0.12.0 library. This clustered heatmap with the dendrogram figure was exported in a vectorized format to the directory “concatenated_scores”.

#### GATOR Conservation Diagram

Single GATOR windows were plotted as individual figures using pyGenomeViz v0.4.4 (41) along with labels to provide genomic context information. These labels consisted of the track name of the GATOR window, protein annotations for required and optional genes, and the genomic length along with its scale. Genes were color-coded based on the user’s query type: required genes in purple, and optional genes in orange. Genes not included in the protein queries were colored green. The edges of genes located at the contig edge were highlighted in red. The percentage of conservation of each gene across the GATOR windows was computed from the gene-level presence-absence tables. Thus, the percentage of conservation for each gene was illustrated by adjusting the transparency of the gene colors. Genes that were completely present across all windows (i.e., 100% conserved) were shown as fully opaque, while those completely absent (i.e., 0% conserved) were displayed as fully transparent. Additionally, a heatmap was generated, with varying degrees of transparency to represent the percentage of conservation. These GATOR conservation figures were exported in a vectorized format to the directory named “gator_conservation_plots”.

#### GATOR Focal Neighborhood

All GATOR windows were plotted into single figures, using the same library and genomic context information labels as in the GATOR conservation diagram sub-section. However, protein annotations were displayed only for the first track, which consisted of the GFW. The organization of the subsequent GATOR windows in the plot was determined by the GFS values, with the window having the highest GFS placed directly below the GFW, and so on in descending order with the lowest GFS positioned at the bottom. Gene homology rails were incorporated based on the protein alignment positions from the output table generated by DIAMOND. For modular genes, the homology rail represented the hit with the highest bit score. In contrast, for non-modular genes, all homology rail hits were plotted. Therefore, each GFW featured a unique arrangement of GATOR windows based on their similarities. These GATOR neighborhood figures were then exported in a vectorized format to the directory “gator_neighborhood_plots”.

#### Targeted GATOR-GC Database

All genomes from AllTheBacteria release 0.1 (∼1.92M; (42)) and the Natural Products Discovery Center (NPDC) portal (8k; (43)), and all bacterial BGCs (∼2k) from the MIBiG v3.0 database were downloaded in FASTA format as of October 2023. These FASTA files were used to train and predict genes using Prodigal v2.6.3 (44). Feature qualifiers for the predicted genes and amino acid sequences were outputted in GenBank format files. To further explore Actinomycetes genomes, another targeted database was created, including the NPDC genomes, the Actinomycetes genomes (∼55k) from NCBI (retrieved June 2024), and all bacterial MIBiG v3.0 BGCs. These two sets served as input for PRE-GATOR-GC to generate the DIAMOND and modular domains databases. The taxonomy classification for these databases was performed using classify_wf in the GTDB-Tk v2.3.0 (45).

### MIBiG Global Analysis

#### BGC Identification

The bacterial MIBiG BGC dataset was converted into FASTA DNA format files, which served as queries for antiSMASH v7.0. The relaxed strictness parameter was used for the HMM-based cluster detection method. The Prodigal algorithm and a minimum contig length of 1kb were specified for the BGC identification process. Furthermore, additional analyses within antiSMASH were conducted, including comparisons of identified clusters against a database of antiSMASH-predicted clusters, exploration of known subclusters responsible for synthesizing precursors, and referencing known gene clusters from the MIBiG database.

#### Manual Curation for Overlooked BGCs

Functional annotations for CDSs within unidentified BGCs were manually curated to identify both required and optional proteins needed for the biosynthesis of their products. Fragmented MIBiG BGCs or those with only one gene, duplicate MIBiG BGCs, and those whose publication(s) did not contain the described BGC were not considered for subsequent curation. Selection criteria for required proteins included gene knockout or overexpression experiments that either enhanced or abolished biosynthesis, gene-level expression analysis, gene complementation assays, and gene conservation across similar pathways. Optional proteins were chosen based on their role in adding structural diversity to the final product without hindering its production, such as tailoring enzymes or transport enzymes. For each MIBiG BGC, protein sequences, and their identifiers were systematically extracted to generate the required and optional FASTA protein query files to run GATOR-GC.

#### Targeted Genome Mining

All required and optional proteins identified above were combined into a single FASTA query file. This file was aligned against the targeted GATOR-GC database using DIAMOND with a block size of 20, and a chunk size of 1.0. This DIAMOND output was parsed by GATOR-GC using the required and optional proteins for each MIBiG BGC as input. The default parameters were used for HMMER’s hmmsearch in the detection of modular domains. The required intergenic distance and window extension parameters of GATOR-GC were also set to their default values.

#### Network Analysis of GATOR Windows of Overlooked MIBiG BGCs

A network displaying the relationships of GATOR windows was created. GFS values calculated for the unique GATOR windows were used to create an edge table that displays the connections between GATOR windows. A node table was created, including metadata such as biosynthetic classes, MIBIG BGC identifiers, and GTDB-TK taxonomy. Gephi toolkit v0.1.1 (46) was used to construct the network using the Force Atlas 2 algorithm, configured with stronger gravity settings, a tolerance speed value of 1.0, 10, 000 iterations, and a scaling factor of 9.5. The group of GATOR windows connected to a specific MIBiG BGC is referred to as a BGC family.

#### GATOR Window Diversity Estimation

Shannon Entropy (H) was calculated to assess the diversity of GATOR windows across different taxonomic levels (from phylum to species). For each BGC family, the number of GATOR windows at each taxonomic level was first counted. These counts were then divided by the total number of GATOR windows in the BGC family. These relative abundances represented the probability of randomly selecting a GATOR Window from a given taxonomic level within the BGC family. The H formula,

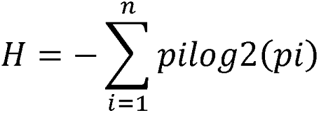

was then applied, where represents the relative abundance of each taxonomic level, and is the total number of taxonomic levels in the BGC family.

H values were categorized based on the interquartile range (IQR) of all entropies. Values within the IQR were called low, those spanning from the upper quartile to the right whisker were called medium, and values exceeding the right whisker were called high.

### Use Case Analysis

Protein sequences for required and optional queries used to call gene clusters for mycosporine-like amino acids (MAAs), 7-deazapurine in DNA (dpd) genomic islands, and FK-family were retrieved from the NCBI (Supplementary Table S1). Taxonomic classification of GATOR windows was performed using classify_wf in GTDB-Tk v2.3.0. MAA and dpd gene clusters were searched within the full targeted database (ATB, NPDC, and MIBiG), while FK-family gene clusters were queried in a subset of the targeted database (Actinomycetes NCBI, NPDC, and MIBiG). The GATOR windows protein database was analyzed using HMM profiles from the PFAM database (47), employing the hmmsearch algorithm in HMMER v3.3.2 to identify protein domains.

MAA gene clusters were called using default GATOR-GC parameters, including protein query coverage, protein identity, required distance, and window extension. Hits for required proteins within unique MAA GATOR windows were aligned and concatenated using MAFFT v7.515 (48). A multilocus phylogeny was then constructed in IQ-TREE v2.1 (49) using Maximum Likelihood inference, applying the best-fit evolutionary model, and performing 100 bootstrap replicates. The resulting consensus tree was edited in iTOL v6 (50), with branch lengths omitted to simplify the visualization.

dpd genomic islands were identified using ultra-sensitive alignment with DIAMOND v2.6.1, applying default parameters for the required distance and a 20kb window extension. A heatmap was constructed using the conservation percentages of *dpd* genes at the phylum level. The heatmap was clustered using the complete linkage method, based on the presence-absence matrix of *dpd* genes. To visualize genomic architecture, GATOR window representatives were selected for each phylum (Supplementary Table S2).

FK-family gene clusters were called using default parameters for DIAMOND (e.g., protein query coverage, and protein identity), a required distance of 15kb, and a window extension of 25kb. GATOR windows with similar genomic architectures were identified using the clustered heatmap on the GFS values automatically generated by GATOR-GC. To visualize their genomic architectures, GATOR window representatives were selected for each FK-family BGC class (Supplementary Table S2).

## Results

### Overview of the GATOR-GC algorithm for identification and comparative analysis of gene clusters

We developed GATOR-GC, a Python-based algorithm designed to identify and compare GATOR windows (i.e., gene clusters) in a targeted, and flexible manner, allowing users to define required and optional proteins. In a single execution, GATOR-GC compares all identified genomic windows to provide insights into their diversity, deduplicates identical GATOR windows to avoid unnecessary calculations, and generates informative, vectorized figures (Figure 1). Before running GATOR-GC, the PRE-GATOR-GC module creates DIAMOND and modular domain databases from user-supplied input genomes in GenBank format.

**Figure 1.**
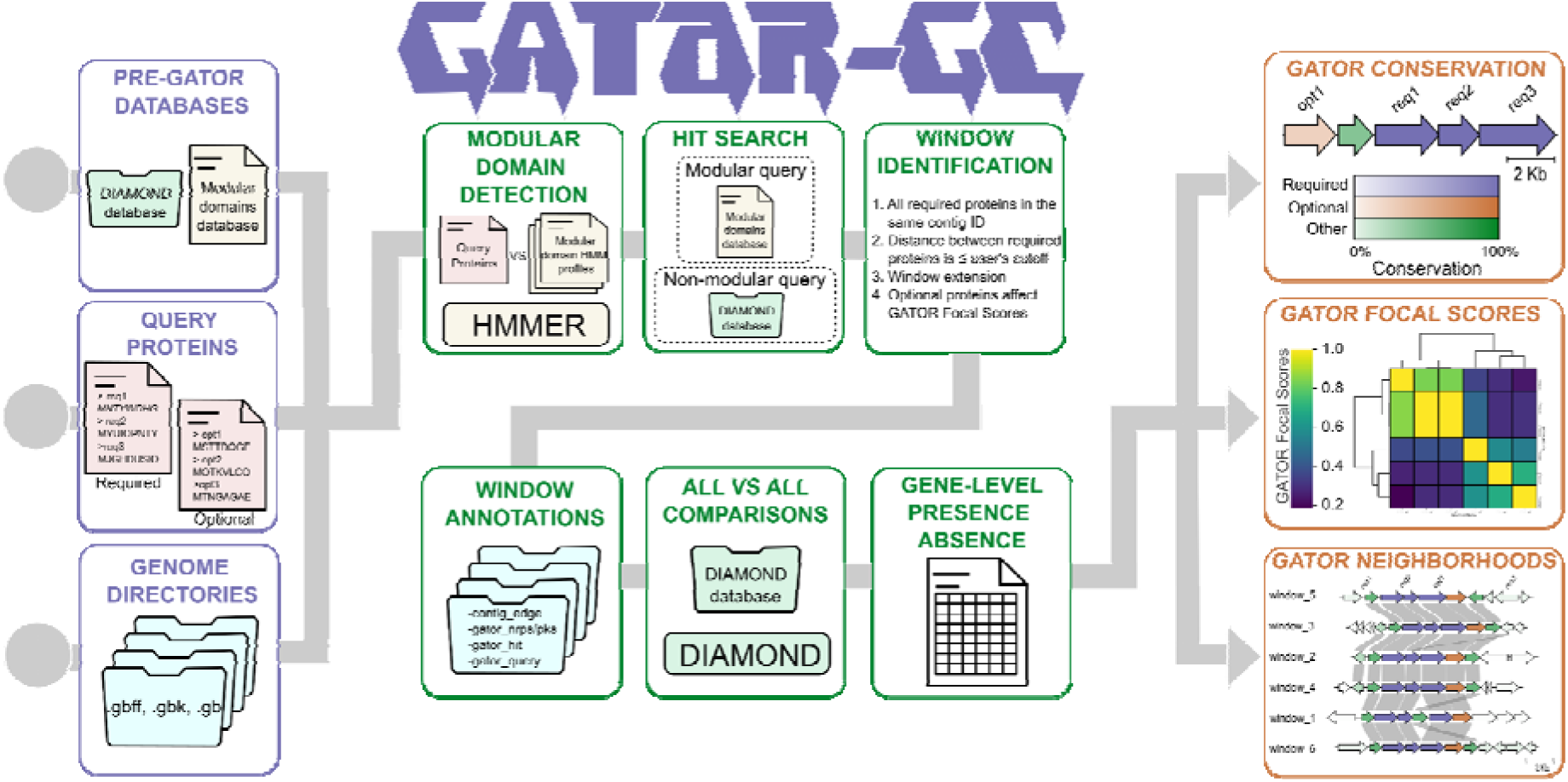
GATOR-GC workflow for gene cluster identification and comparative analysis. The workflow begins with specifying the input files (purple boxes), which include a DIAMOND database, a modular domains database, FASTA protein queries (required and/or optional), and genome sequences in GenBank format. In the processing stage (green boxes), GATOR-GC first screens the query proteins to determine whether modular proteins are present. If modular proteins are detected, their hits are retrieved from the modular domains database. Non-modular query proteins are then searched against the DIAMOND database using DIAMOND to identify hits. Next, these hits are grouped by contigs, and their intergenic distances are evaluated. GATOR-GC calls gene cluster windows if all required proteins are both i) found on the same contig and ii) within the user-defined intergenic distance. While optional genes and window extensions do not influence window identification, they affect GATOR Focal Score (GFS) calculations. GATOR-GC then generates annotated GenBank files for each window and builds a window protein database for all-vs-all comparisons. It creates gene-level presence-absence tables and calculates GFS values for proximity-weighted similarity scoring. To avoid unnecessary calculations, GATOR-GC performs deduplication of identical windows. Outputs (orange boxes) include gene cluster conservation diagrams, a clustered heatmap of GFS, and gene cluster neighborhood diagrams sorted by GFS.

#### Input files

To execute GATOR-GC, users provide the path(s) to the created databases with PRE-GATOR-GC, the genomes of interest, and the FASTA files containing the query proteins.

Required queries, those proteins essential for defining the gene cluster, are supplied by the user. These proteins may include core enzymes responsible for the biosynthesis of a particular molecular feature or serve a key functional role within the gene cluster.

Optional queries serve as flexible markers for proteins that may not be ideal for defining a cluster but whose presence may otherwise be of interest. For instance, in secondary metabolism, using methyltransferases or cytochromes P450 to define a BGC would lead to many spurious identifications due to their widespread genomic presence in other contexts. However, when treated as optional queries, their presence within a gene cluster may suggest structural diversity for its product.

This flexible approach to protein queries allows users to test hypotheses regarding the presence or absence of specific proteins within gene clusters. It also facilitates the exploration of how certain genes influence genomic architecture, providing valuable insights into the diversity and conservation of these pathways.

#### Processing Steps

GATOR-GC includes an automatic check for the presence of modular domains within user queries. Hits for modular and non-modular query proteins are stored in a protein dictionary for subsequent GATOR window detection. GATOR-GC identifies a GATOR window only when all required proteins are present on the same contig and within the user-defined intergenic distances. To improve window border predictions, users can adjust the GATOR window size by specifying a genomic length to extend both upstream and downstream regions instead of relying on static gene cluster flank extensions. While this window extension does not affect the identification of GATOR windows, it can influence the GATOR Focal Score (GFS) calculations (as more genes may be included in the window) and can help define GATOR window boundaries.

GFS is a novel metric designed to rapidly evaluate the (dis)similarity of GATOR windows. GFS weights the gene-level presence-absence based on proximity to user-defined queries (i.e., required and/or optional). This proximity-weighted similarity scoring provides a fast and quantitative assessment of window diversity and conservation. Unlike other empirical similarity scoring methods that rely on static, syntenic weighting factors, the GFS system dynamically calculates specific weights for each focal window. These weights depend on the number of genes and their positions relative to the user’s query genes, ensuring a context-specific and dynamic similarity evaluation for genomic synteny (Supplementary Figure S2).

#### Output Files

GATOR-GC generates a summary table of key information for each query protein in a GATOR window. This includes the query type (i.e., required/optional), a boolean classification indicating whether the protein was categorized as NRPS and/or PKS, and genomic context information (i.e., start position, end position, locus ID, contig ID). Additionally, the table specifies the GATOR window ID to which the protein belongs. GATOR window GenBank files are annotated with new feature qualifiers for each CDS. These qualifiers include boolean indicators for whether a CDS is located at the contig edge and whether it is categorized as NRPS and/or PKS. Additionally, the qualifiers specify whether CDS matches the protein user’s query and provide the respective window ID. Gene-level presence-absence and the GFS tables for each GATOR window are also provided. GATOR-GC creates vectorized figures, summarizing the hierarchical clustering of GFS values in a heatmap, alongside gene conservation and genomic neighborhood information. The clustered heatmap with GFS provides insights into the global diversity of all-vs-all GATOR windows and enables the identification of groups with similar genomic architectures. The GATOR gene conservation diagram provides a visualization for each GATOR window where users can identify highly conserved genes, even if they were not included in the query. This can help users refine their query to include proteins that may not have been considered important or conserved. GATOR neighborhood visualization figures include all GATOR windows to display similarity between gene content, following the same color and conservation as the GATOR conservation diagrams. These GATOR windows are sorted by GFS, enabling users to view the most similar windows at the top and the most dissimilar ones at the bottom.

In the following sections, we will showcase the versatility of GATOR-GC in locating and comparing different types of gene clusters and demonstrate its applications in estimating cluster diversity, calculating gene conservation, and identifying distinct genomic architectures.

### GATOR-GC identifies over 4 million GATOR windows related to overlooked MIBiG BGCs

We assessed how frequently bacterial MIBiG v3.0 BGCs (n=1, 999) could be identified using antiSMASH v7.0 (relaxed strictness). In total, 279 BGCs (∼14%) were not detected using this tool (Figure 2A). The genera with the highest proportions of missed BGCs were *Streptomyces* (30.1%), *Pseudomonas* (7.16%), *Escherichia* (5.73%), *Streptococcus* (3.94%), and *Burkholderia* (3.22%). In all genera but *Streptomyces*, the proportion of missed BGCs exceeded those identified. These other genera represent 52.3% and 49.8% of antiSMASH-identified and -missed BGCs, respectively (Figure 2B). We found that modular BGCs (e.g., NRPS, PKS), hybrids (i.e., more than one class), and RiPPs had higher proportions of antiSMASH-identified BGCs than missed ones, indicating antiSMASH’s proficiency in these classes. However, BGCs classified as “other” (i.e., terpenes, alkaloids, and those that do not fit into any other category), and saccharide classes had the highest proportions of missed BGCs and were missed more often than not (Figure 2C).

**Figure 2.**
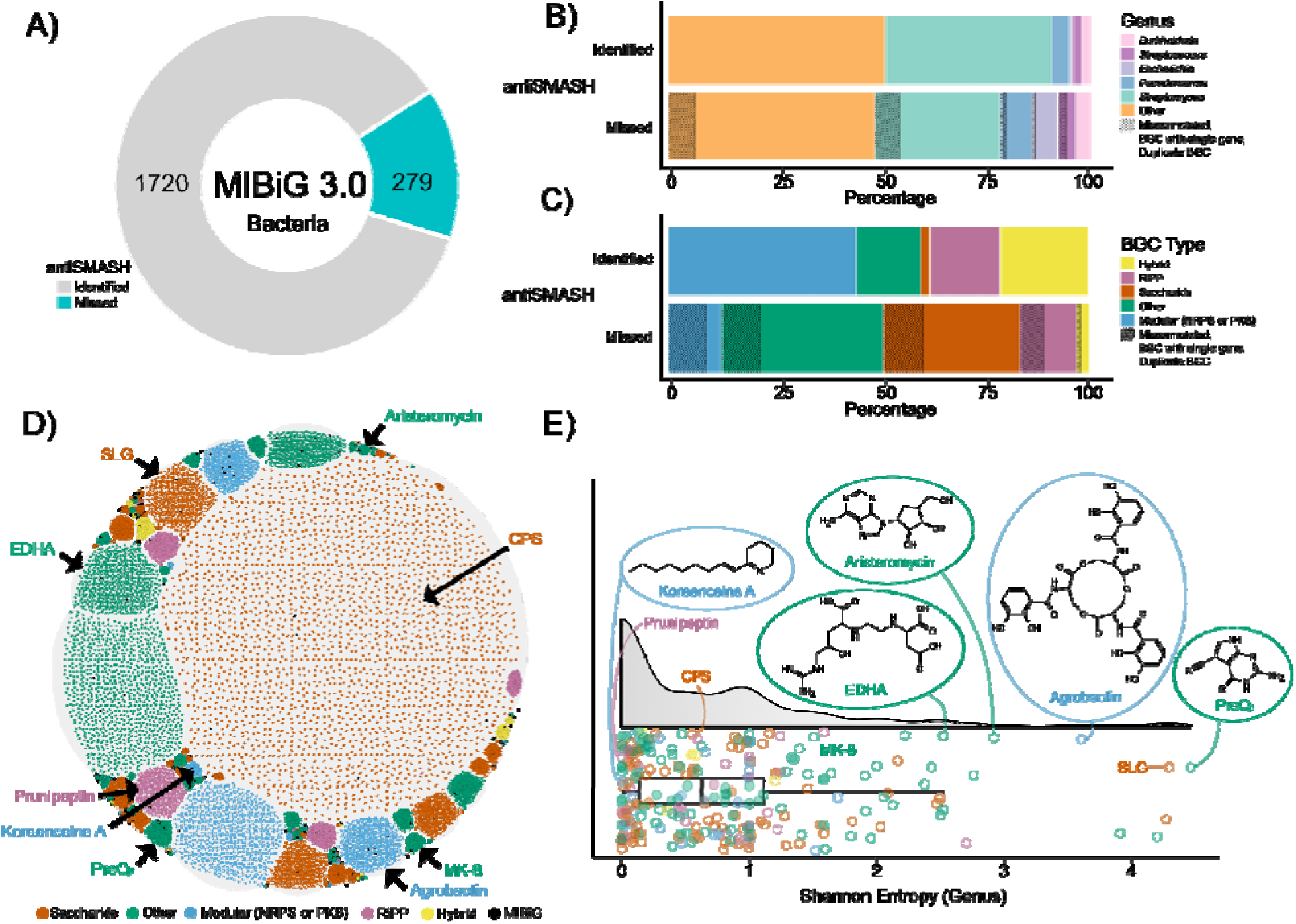
GATOR-GC identified over 4 million related GATOR windows to MIBiG BGCs undetected by antiSMASH, forming 233 BGC families. A) Bacterial MIBiG dataset (n=1, 999) showing the number of BGCs that are identified (n=1, 720) and missed (n=279) by antiSMASH. (B) The proportion of identified and missed BGCs categorized by the MIBiG genus taxonomy and (C) MIBiG BGC types are shown. The black texture in (B) and (C) represents BGCs with problematic annotations (see Methods). D) GATOR window landscape, with each unique GATOR window (n=16, 102) as nodes and GFS as edge values. Nodes are colored according to the MIBiG BGC type, with MIBiG BGCs in black. Each subnetwork corresponds to a different BGC family. Black arrows indicate subnetworks highlighted in (E). E) Distribution of genus-level Shannon entropies for each BGC family, colored according to the MIBiG BGC types. Highlighted examples from (D) include their respective MIBiG BGC products and/or their chemical structures. CPS: capsular polysaccharide; EDHA: ethylenediaminesuccinic acid hydroxyarginine; SLG: S-layer glycan.

We manually verified the reported metadata and the experimental validation of these BGCs. Our curation revealed issues in some of these BGCs, including duplicate entries, single-gene BGCs, discrepancies between the MIBiG biosynthetic classes and the biosynthesis types described in their publication(s) (e.g., polyketide pathways lacking a PKS), and BGCs that have only a flank region of the full biosynthetic pathway. As a result, 46 of the 279 missed BGCs were categorized as problematic annotations (proportions indicated with black texture in Figure 2B and Figure 2C) and removed, leaving a refined dataset of 233 BGCs for downstream analysis.

To identify these 233 missed BGCs, we used GATOR-GC to detect similar gene clusters across ∼1.92M publicly available genomes. We manually classified proteins as required or optional according to their biosynthetic role(s) as described in their source publications (see MIBiG Global Analysis, Manual Curation for Overlooked BGCs section in Methods) (Supplementary Table S3). We ran GATOR-GC independently for each BGC and identified 4, 201, 1875 total and 16, 102 unique GATOR windows similar to the missed MIBiG BGCs. Of the 233 BGC families, 16 were singletons (i.e., only the MIBiG BGC was identified). BGC family members encoding saccharides comprised 50.15% of identified windows, with “other” representing 30.51%, modular 11.15%, RiPPs 6.97%, and hybrids 1.22% (Supplementary Figure S3). These GATOR windows spanned different taxonomic groups. The genera contributing the most to these BGC families were *Klebsiella* (25.84%), *Streptomyces* (23.75%), *Escherichia* (8.29%), *Pseudomonas* (4.81%), *Mycobacterium* (3.10%), *Burkholderia* (2.04%), and *Streptococcus* (1.75%) (Supplementary Figure S4A).

To explore the relationships within BGC families, we constructed a multi-network graph using the GATOR’s GFS values for the deduplicated (i.e., unique) BGCs (nodes=BGCs; edges=GFS). These BGC families formed discrete densely connected sub-networks, with a total of 6, 891, 949 connections, reflecting the similarity between GATOR windows within the same BGC family (Figure 2D). We observed that most BGC families were composed of GATOR windows from bacterial genomes of multiple genera (Supplementary Figure S4B).

We calculated Shannon Entropy (H) to quantify genus-level diversity within each BGC family (Supplementary Table S4). Of the 233 BGC families, 179 (76.82%) were classified with low H (≥0 and ≤1.2), 45 BGC families (19.31%) had medium H (>1.2 and ≤2.6), and 9 BGC families (3.87%) had high H (>2.6) (Figure 2E). To provide some taxonomic context for these BGC families, we highlighted representative examples for each H level (Figures 2D and 2E).

BGC families with low H included those encoding for koreenceine A (MIBiG accession BGC0001987), the graspetide prunipetin (BGC0002725), and capsular polysaccharide (BGC0000728). The koreenceine BGC family comprised 664 total members, predominantly represented by *Pseudomonas* (99.39%, n=654) and spanning five genera. The prunipeptin BGC family included 3, 589 total members distributed across six genera, with *Streptomyces* having the largest representation (99.53%, n=3, 572). The capsular polysaccharide BGC family had 56, 715 total members across 31 genera, with *Klebsiella* contributing 89.16% (n=50, 565).

BGC families with medium H included those encoding menaquinone (MK-8, BGC0000913) and ethylenediaminesuccinic acid hydroxyarginine (EDHA, BGC0002567). The menaquinone BGC family had the largest number of total windows in the dataset, with 943, 199 total members distributed across 115 genera. This BGC family was primarily represented by *Salmonella* (57.68%, n=544, 072), and *Escherichia* (33.03 %, n=311, 578). The EDHA BGC family comprised 1, 450 total windows spanning 24 genera, with contributions from *Streptomyces* (41.17%, n=597), *Nocardia* (22.62%, n=328), and *Corynebacterium* (18.34%, n=266).

High H was observed in BGC families encoding aristeromycin (BGC0001514), agrobactin (BGC0002474), S-layer glycan (BGC0000794), and 7-cyano-7-deazaguanine (preQ_0_, BGC0001217). The aristeromycin BGC family included 10 total members evenly distributed across six genera, such as *Saccharopolyspora* (20%, n=2), *Streptomyces* (10%, n=1), *Actinoplanes* (10%, n=1), and *Micromonospora* (10%, n=1). The agrobactin BGC family contained 20, 596 total windows across 128 genera, with major contributions from *Escherichia* (27.09%, n=5, 579), *Klebsiella* (18.48%, n=3, 807), and *Bacillus* (15.73%, n=3, 239). The S-layer glycan BGC family involved 886 total members across 77 genera, including *Paenibacillus* (33.97%, n=301), *Mycobacterium* (17.04%, n=151), and *Clostridium* (15.12%, n=134). The preQ_0_ BGC family exhibited the highest genus-level diversity, with 828 total windows spanning 95 genera; major contributors included *Streptomyces* (28.02%, n=232), *Nitratidesulfovibrio* (10.87%, n=90), and *Chromobacterium* (8.82%, n=73).

These findings underscore the utility of GATOR-GC in identifying overlooked gene clusters without bias towards biosynthetic classes or taxonomy groups, and without relying on pre-defined biosynthetic rules. In the following sections, we will target various gene clusters to demonstrate additional features of GATOR-GC that may prove valuable to the community.

### Discovery of two previously undescribed conserved genes in mycosporine-like amino acid (MAA) gene clusters

We used GATOR-GC to evaluate the gene-level conservation of gene clusters that encode mycosporine-like amino acids (MAAs), water-soluble secondary metabolites with UV-protectant properties found across eukaryotic and prokaryotic organisms. All known MAAs have in common a six-membered carbon ring, known as the cyclohexanone or cyclohexenimine scaffold (51). MAAs have gained increasing attention due to their potential to be used as sunscreen ingredients. Thus, identifying novel BGC architectures with unknown enzymes could serve as a starting point for the elucidation and characterization of potential new MAA analogs with commercial potential.

The minimum set of enzymes necessary to produce a cyclohexanone scaffold was used as required proteins for GATOR-GC, namely a dehydroquinate synthase (DHQS), an *O*-methyltransferase (O-MT), and an adenosine triphosphate (ATP)-grasp domain-containing protein (52). As optional query proteins, we selected the enzymes whose participation contributes to the structural diversity of MAAs. These optional proteins included an NRPS-like enzyme, a D-alanyl-D-alanine ligase, a phytanoyl CoA dioxygenase (53), and a short-chain dehydrogenase/reductase (SDR; (54)).

With GATOR-GC we identified 208 total and 131 unique GATOR windows across different bacterial phyla, including Armatimonadota, Bacillota, Planctomycetota, Pseudomonadata, Cyanobacteriota, and Actinomycetota. We also identified the MIBiG BGCs (BGC0000427 and BGC0001615), which encode the biosynthesis of 4-deoxygadusol, mycosporine glycine, shinorine, hexose-palythine-serine, and hexose-shinorine (Supplementary Table S5).

To evaluate the presence of additional proteins that may be related to the biosynthesis of MAAs, we used the GATOR gene conservation feature to select the GATOR window with the highest percentage of conserved proteins: a window from *Pseudonocardia alni* (Assembly SAMN14595976) that has two well conserved (> 60%) co-occurring genes (Figure 3A). We identified their domains using HMM profiles in the PFAM database. These domains were associated with haloacid dehalogenase (HAD) and taurine catabolism dioxygenase (TauD/TfdA), which belongs to the broader nonheme Fe(II) 2-ketoglutarate dependent superfamily of dioxygenases (e.g., including phytanoyl CoA dioxygenases) (Figure 3B). To the best of our knowledge, there are no known BGCs that code for MAAs harboring HAD or TauD/TfdA. This finding expands the known enzymatic diversity associated with these BGCs and opens opportunities for novel MAA analog discovery.

**Figure 3.**
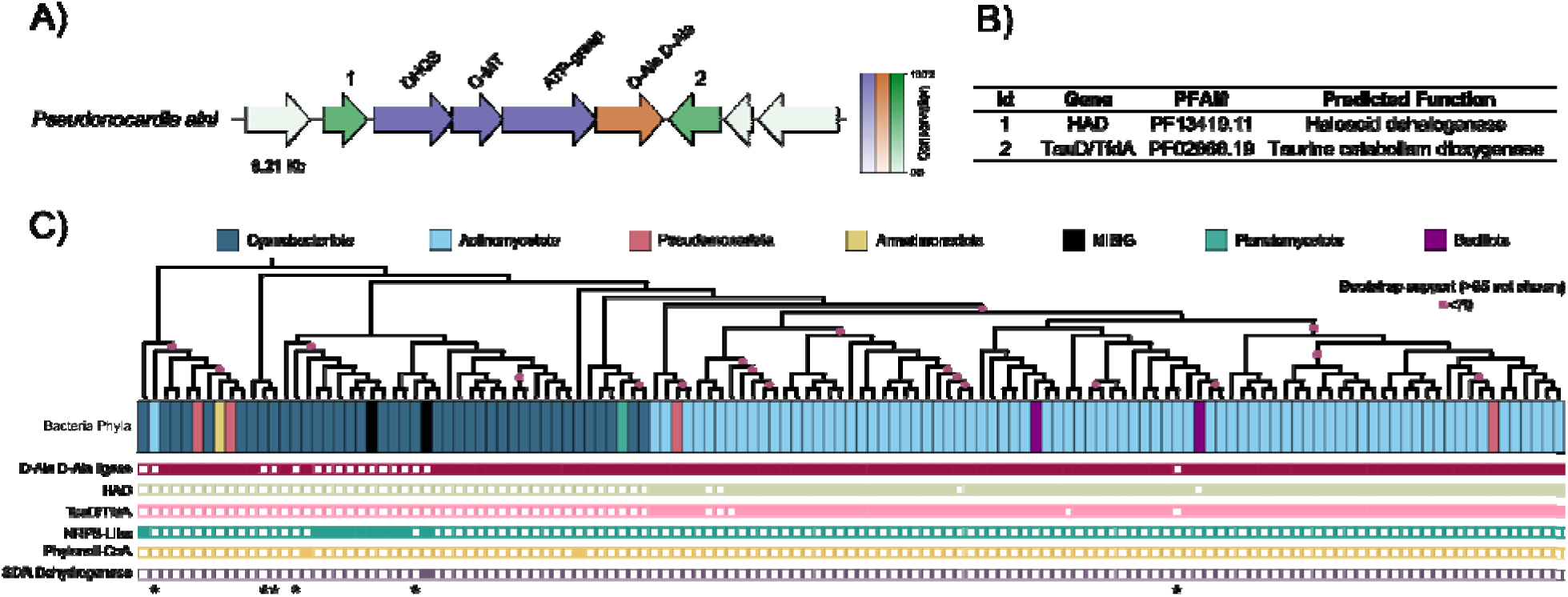
Conservation and phylogenetic analysis of enzymes involved in the biosynthesis of MAAs. A) GATOR-GC conservation figure for a putative MAA BGC of *Pseudonocardia alni* showing enzyme conservation across the dataset. Genes colored in purple represent the required proteins (DHQS – dehydroquinate synthase, O-MT – *O*-methyltransferase, ATP-grasp – adenosine triphosphate (ATP)-grasp superfamily). Genes in orange represent optional enzymes (D-Ala-D-Ala – D-alanyl-D-alanine ligase). Genes in green depict other proteins not used in the GATOR-GC search. The color bar indicates the degree of conservation for each gene in the BGC across the dataset. Genes colored green with numbers at the top of the arrows (1 and 2) represent other genes that are well conserved in the dataset. B) Predicted PFAMs and biochemical functions for the conserved genes identified by GATOR-GC (1 – HAD, 2 – TauD/TfdA). C) Phylogenetic distribution constructed with required proteins of 131 putative MAA BGCs along with the presence-absence of enzymes involved in their biosynthesis at the bottom, and the GTDB-Tk taxonomy at the phylum level at the top. Bootstrap support values below 70 % are shown. Asterisks (*) at the bottom of the enzyme-presence-absence matrix represent putative BGCs that do not have either an NRPS-like or a D-Ala-D-Ala ligase, suggesting the formation of cyclohexanone scaffold.

To map the taxonomic distribution of the biosynthetic enzymes involved in the biosynthesis of MAAs, including the conserved proteins (HAD, and TauD/TfdA), we built a multilocus phylogeny using the required proteins from the 131 unique windows. We found that NRPS-like enzymes were present only in a subset of Cyanobacteriota, while the D-alanyl-D-alanine ligase was more widespread. HAD and TauD/TfdA were not present in Cyanobacteriota but were found in Actinomycetota, Pseudomonadata, and Bacillota (Figure 3C).

A total of six windows, five in Cyanobacteriota and one in Actinomycetota did not have any hits for the optional proteins. We analyzed protein-encoding genes within these windows with HMM profiles of the PFAM database and found some protein domains associated with metallopeptidases, glycosyl hydrolases, aminobenzoate oxygenase, guanylate cyclase, and some domains of unknown function (DUF389) (Supplementary Figure S5). The full list of unique GATOR windows, the species-level taxonomy, and the presence-absence data for the proteins (required/optional, HAD and TauD/TfdA) involved in MAA biosynthesis is reported in Supplementary Table S5.

In summary, GATOR-GC identified two additional proteins that are highly conserved in MAA gene clusters. Although not previously linked to MAA biosynthesis, their consistent co-occurrence suggests a potential role in expanding MAA diversity. Here, GATOR-GC improved the delineation of gene cluster boundaries beyond the initial search criteria by evaluating the conservation of genes in an evolutionary context.

### GATOR-GC identified taxonomic and evolutionary patterns of 7-deazapurine in DNA (*dpd)* genomic islands across bacteria

Next, we aimed to identify dpd genomic islands that modify DNA with 7-deazapurine derivatives across bacteria. 7-deazapurines are nucleoside analogs derived from 7-cyano-deazaguanine (PreQ_0_), the universal precursor of all known 7-deazapurine-containing molecules (55). PreQ_0_ biosynthesis occurs in a four-step pathway from guanosine-5’-triphosphate (GTP). First, GTP cyclohydrolase I (FolE) catalyzes the conversion of GTP into 7, 8-dihydroneopterin triphosphate (H_2_NTP). Next, 6-carboxy-5, 6, 7, 8-tetrahydropterin synthase (QueD) converts H_2_NTP into 6-carboxy-5, 6, 7, 8-tetrahydropterin (CPH4). The third step involves the radical SAM enzyme 7-carboxy-7-deazaguanine synthase (QueE), which converts CPH4 into 7-carboxy-7-deazaguanine (CDG). Finally, 7-cyano-7-deazaguanine synthase (QueC) catalyzes the conversion of CDG into preQ_0_ (56).

7-deazapurines play multifaceted roles in bacteria, they can serve as scaffolds for secondary metabolite biosynthesis and contribute to restriction-modification systems when incorporated into nucleic acids (56). For example, *Salmonella enterica* serovar Montevideo (*S. Montevideo*), *Kineococcus radiotolerans*, *Comamonas testosteroni,* and other bacteria harbor genomic islands that modify DNA with 2’-deoxy-preQ_0_ (dPreQ_0_) and 2’-deoxy-7-amido-7-deazaguanosine (dADG) (55). In *S. Montevideo,* these modifications inhibit the replication of unmodified DNA, suggesting a self/non-self discrimination mechanism (55).

The dpd genomic islands consist of up to eleven genes organized as a synthesis operon (*dpdA-C*) and a restriction operon (*dpdD*-*dpdK*) (57). Since these islands are widespread in bacteria and exhibit variable gene content, GATOR-GC is ideal for their identification given its flexibility to capture evolutionary diversity.

To identify these dpd islands, we set *dpdAB* as required and *dpdC* along with the restriction operon as optional proteins. Although *dpdC* is required for dADG formation in *S. Montevideo* (57, 58), not all experimentally validated dpd islands encode dADG (55). Therefore, setting *dpdC* as optional allows us to capture a broader diversity of islands, including those that may only produce dPreQ . GATOR-GC analysis yielded 37, 198 total and 2, 335 unique windows in eight bacterial phyla and 118 genera (Supplementary Table S6).

To investigate the genomic architectures and explore putative taxonomic and evolutionary patterns of these unique windows, we calculated the percentages of conservation of *dpd* genes at the phylum level. The hierarchical clustering-based dendrogram on the conservation values revealed two main groups, each comprising four phyla with similar *dpd* gene contents (Figure 4A). The first contained two sister subgroups: one clustering Bacillota (n=11) with Actinomycetota (n=333) and another grouping Pseudomonadota (n= 1, 971) with Bacteriodota (n=3). This group was characterized by the presence of the complete synthesis and restriction operons. The second group consisted of a single subgroup where Verrucomicrobiota (n=1) and Myxoccota (n=6) clustered together. Campylobacterota (n=7) formed a sister branch to this cluster, and Cyanobacteriota (n=1) was a separate sister branch encompassing all members of the second group. This group exhibited reduced restriction operon content, with some restriction genes retained in Myxococota and Campylobacterota, whereas Verrucomicrobiota and Cyanobacteriota lacked all restriction genes.

**Figure 4.**
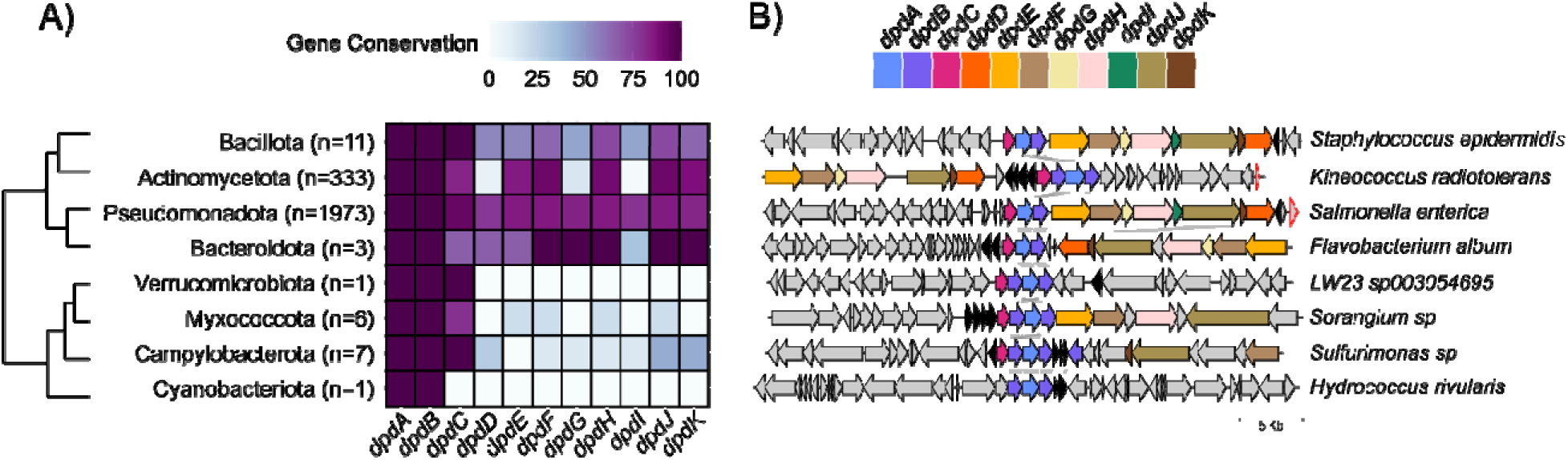
Gene conservation across 2, 335 unique *dpd* genomic islands. A) Heatmap illustrating the conservation of *dpd* genes across bacterial phyla, with the number of unique genomic islands shown in parentheses. Color intensity represents the percentage conservation of each gene, with darker shades indicating higher conservation. The dendrogram was constructed using the complete linkage method, based on the presence-absence matrix of *dpd* genes generated by GATOR-GC. B) Representative GATOR neighborhoods of *dpd* genomic architectures are shown for each phylum, highlighting the structural variability and conservation patterns within these genomic islands. Genes colored black represent proteins associated with the biosynthesis (FolE, QueE, QueD, and QueC) and transport (YhhQ) of preQ_0_.

The identified genomic islands were predominantly associated with genera such as *Escherichia, Klebsiella*, *Salmonella*, *Streptomyces*, *Pseudomonas*, *Enterobacter*, and *Citrobacter*, which collectively accounted for 80.78% of the total (Supplementary Figure S6A). The remaining 111 genera were less represented, each contributing fewer than 40 windows (Supplementary Figure S6B). The clustered heatmap at the genus level revealed a complex grouping pattern, primarily influenced by the number of genera and the predominant occurrence of single windows (Supplementary Figure S6C). Genomic islands lacking the restriction operon were divided into two groups: those with *dpdC* (e.g., *Aliarcobacter*, *Agrobacterium*, *Archangium*) and those without *dpdC* (e.g., *Croceibacterium*, *Acidovorax*, and *Ferrovum*).

Overall, 146 distinct *dpd* gene compositions within these islands were identified, with fully intact operons being the most common (>50%), while those lacking some genes were less frequent (Supplementary Table S7). Most *dpd* genes clustered in over 70% of the *dpd* islands with *dpdAB*, except for *dpdD*, and *dpdI*. Notably, *dpdC* was the most frequently clustered gene with *dpdAB*. Multiple copies of many *dpd* genes were observed. Among them, *dpdB* exhibited the highest number of additional copies (n=801), while *dpdK* had only a single additional copy (Supplementary Table S8).

To evaluate the genomic context of these genomic islands, we analyzed their protein-encoding genes using HMM profiles from the PFAM database (Supplementary Table S9). Frequently detected domains were linked to putative DNA-binding enzymes, including DEAD/DEAH box helicase family (e.g., *dpdFJ*), DNA-sulfur modification-associated domains (DndB, e.g., dpdB), queuine tRNA-ribosyltransferases (TGT, e.g., dpdA), PLD-like domains (e.g., dpdK), ATPase domains involved in DNA replication (e.g., dpdH), SNF2-type helicases (e.g., dpdE), and a domain of unknown function (DUF6884, e.g., dpdC). Additionally, enzymes frequently co-localizing with *dpd* genes included QueD (PTPS, PF01242), detected in 78% of the islands, as well as DNA-binding domains (Arm, PF13356), zinc-binding dehydrogenase domains (PF00107), QueC (PF06508), lipopolysaccharide export system permeases (PF03739), QueE (PF04055), cytosol aminopeptidases (PF00883), DNA polymerase III chi subunits (PF00133), and transposase-like domains (PF01527), all present in over 20% of the islands.

Some of these enzymes (e.g., FolE, QueE, QueD, and QueC) are involved in the biosynthesis of preQ_0_, a shared precursor for 7-deazapurines (56). Further analysis of representative GATOR windows (Figure 4B) showed strong synteny of the *dpdA-C* genes, although the presence of *dpdD-K* genes varied. Beyond the *dpd* genes, other neighboring genes included those encoding enzymes for preQ_0_ biosynthesis and its YhhQ transporter. Additional flanking genes encoded transposases-like proteins, nucleotide-binding domains, toxin-antitoxin systems (ccdA: PF07362, ccdB: PF01845), and glyoxalase/bleomycin resistant proteins (PF00903). Islands from *LW23* and *Hydrococcus rivularis* lacked the restriction operon, with *H. rivularis* also missing *dpdC*. These islands clustered with other elements, including a phage integrase (PF00589), a glycosyl hydrolase family member (PF01120), His-Asn-His (HNH: PF01844) endonucleases, and several hypothetical proteins. Among these representatives, the presence of 7-deazapurine derivatives in DNA has been experimentally validated in *S. Montevideo* and *Kineococcus radiotolerans* (55).

Collectively, these findings revealed specific distribution patterns of *dpd* genomic islands, highlighting variations in gene content that may influence 7-deazapurine DNA modifications and restriction-modification systems. The flexibility of GATOR-GC, particularly its optional protein query feature, enabled the classification of islands into diverse genomic architectures, potentially revealing novel 7-deazapurines involved in DNA modifications.

### GATOR Focal Scores (GFS) group the FK BGC Family into four distinct classes according to their chemistry

We aimed to evaluate the presence and distribution of the FK BGC family in Actinomycetes. The FK family consists of potent macrolide compounds such as rapamycin, FK520, FK506, and WDB003, representing an important group of secondary metabolites produced by Actinomycetes. These compounds are renowned for their immunosuppressive and antifungal properties. Rapamycin and FK506/520 bind FK506-binding protein 12 (FKBP12) (59) and this complex further targets either the mammalian target of the rapamycin (mTOR) or calcineurin, respectively, making them valuable immunosuppressants that have been used to prevent organ transplant rejection (60). The compound WDB003 (produced by cluster X1) similarly binds FKBP12 and targets the centrosome-associated protein CEP250, which is involved in centrosome cohesion during cell division (61).

To identify members of this family, our required search criteria included the enzymes that synthesize the conserved scaffold at the FKBP-binding interface, such as lysine cyclodeaminase (KCDA), a nonribosomal peptide responsible for pipecolate incorporation (NRPS), a PKS, and a chorismatase. We included optional proteins for an O-MT, a cytochrome P450, and those involved in the biosynthesis of the side chains at C21 specific to FK506 and FK520 (namely, TcsD, TcsC, TscB, and TscA for FK506 and FkbU, FkbE, and FkbS for FK520) (62).

We identified 27 total and 24 unique GATOR windows, along with the MIBiG entries for rapamycin (BGC0001040) and FK520 (BGC0000994). These putative BGCs were mostly found in *Streptomyces* species.

Furthermore, we evaluated the GFS clustered heatmap automatically generated by GATOR-GC to identify similar BGC architectural groups. This all-vs-all comparison analysis revealed that these GATOR windows were grouped into known BGC classes (Figure 5A). These classes corresponded to three different GATOR windows for FK520, five for X1, six for rapamycin, seven for both FK506, and two truncated GATOR windows related to the rapamycin group. These distinctive BGC classes were associated with the known chemical structures for WDB003, FK520, FK506, and rapamycin (Figure 5C). The full set of 24 unique GATOR windows and their neighborhoods is provided in Supplementary Figure S7.

**Figure 5.**
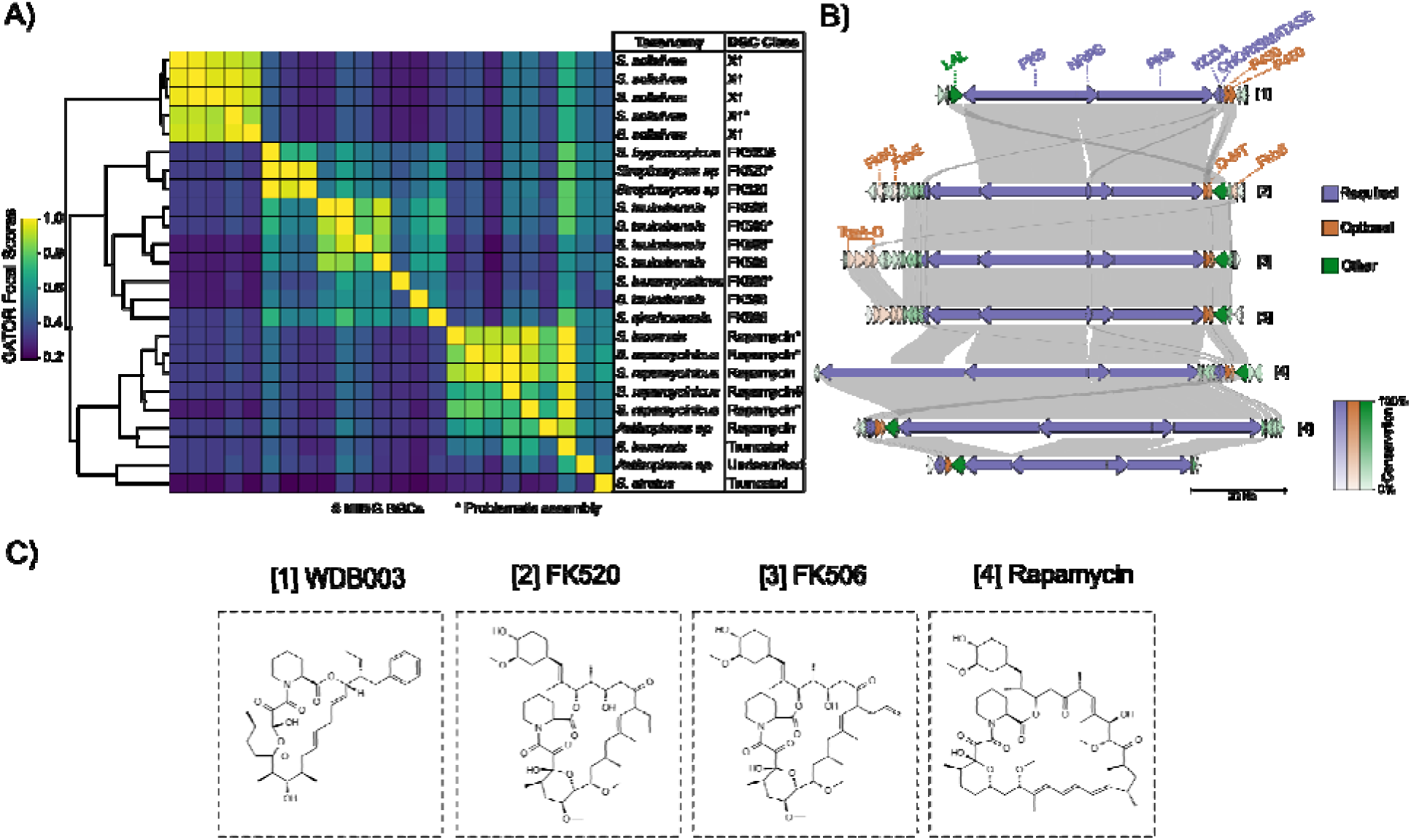
GATOR-GC identified 24 unique GATOR windows belonging to the FK family in Actinomycetes. A) Clustered heatmap of the GFS values generated by GATOR-GC. Each row and column in the heatmap represents a GATOR window. The MIBiG BGC ID, taxonomy at the genus level, and BGC classes are displayed for each GATOR window. BGC classes marked with an asterisk (*) highlight GATOR windows with problematic genomic assemblies while MIBiG BGCs are marked with a #. Truncated BGC classes correspond to GATOR windows found at contig edges. B) GATOR window neighborhoods for representative FK members. Required protein labels are shown in purple, optional in orange, and others in green. The GATOR conservation figure indicates that the LuxR family regulator was well-conserved across the BGC family. C) Chemical structures for (1) WDB003, (2) FK520, (3) FK506, and (4) rapamycin.

The X1 BGC is characterized by an NRPS situated between two PKSs, with KCDA and chorismatase located adjacent to two P450 enzymes (O-MT is absent). In contrast, the FK520 and FK506 BGCs contained the chorismatase and NRPS within three PKSs; however, the KCDA is not adjacent to the P450 and O-MT, but rather upstream of a PKS.

GFS similarity scoring successfully distinguished between GATOR windows related to FK520 and FK506, primarily due to differences in their C21 side chain chemistry: the FK520 BGC contains genes encoding for FkbU, FkbE, and FkbS, which are responsible for the ethyl group at C21, while the FK506 BGC contains genes for TcsA, TcsB, TcsC, and TscD, responsible for the allyl group at the same location. Although the absence of these optional proteins did not prevent the identification of windows lacking them, GFS weighted the genes neighboring these optional proteins more heavily. Additionally, we observed that the FK506 BGC exhibited two distinct gene content organizations. In particular, GATOR windows from *Streptomyces tsukubensis* contained five extra genes adjacent to the TcsA-D region, which were absent in *Streptomyces qinzhouensis*.

The rapamycin BGC featured an NRPS within three PKSs, with chorismatase, KCDA, P450, and O-MT located adjacent to each other. However, we found that a rapamycin BGC found in *Actinoplanes* deviates from the *Streptomyces rapamycinicus* BGC in the positioning of the KCDA, chorismatase, P450, and O-MT, which were upstream of the PKSs rather than downstream. We also identified two BGC architectures in *S. iranensis* and *S. atratus* that were closely grouped with the rapamycin BGC class, though their neighborhoods were truncated (at contig edges). GATOR gene conservation revealed that the LuxR regulator was highly conserved across all windows.

In this case study, we demonstrated the usefulness of GFS in grouping BGCs with similar architectures and mapping them to their known chemistries. GATOR-GC’s ability to identify genomic differences and differentiate between family-related BGCs underscores its potential as a powerful tool for exploring biosynthetic diversity.

## Discussion

GATOR-GC was built to provide a targeted and flexible framework to explore the diversity of gene clusters in a single execution, eliminating the need to chain multiple tools or manual processes. With GATOR-GC, researchers can address several questions, such as i) Are there loci similar to my gene cluster based on required/optional protein search terms, and in which taxonomic groups are they located? ii) Which loci exhibit the greatest syntenic similarity to my gene cluster, and which exhibit the greatest divergence despite containing required genes? iii) What specific genes account for these differences? iv) What is the gene conservation across gene clusters, and are there any genes not used as query search terms whose conservation suggests they may play functional roles? v) Are there sub-families of gene clusters based on similar genomic architectures? By addressing these and other questions, researchers can formulate hypotheses that provide deeper insights into gene cluster diversity. This could help uncover evolutionary patterns in gene cluster distribution and architecture. Moreover, it can aid in the identification of gene clusters overlooked by other genome mining tools. These clusters could play important roles in bacterial fitness or have applications in biotechnology and medicine–roles that may remain unknown unless such clusters are identified and studied.

GATOR-GC uncovered biosynthetic diversity for bacterial MIBiG BGCs that were unable to be detected by antiSMASH. Although antiSMASH is regularly updated to improve its rules and algorithms, not all potential search strategies are implemented in every version. antiSMASH v7.0, used in this study, can annotate 81 different BGC classes (27); however, certain BGCs with unique features or alternative rule sets often go undetected and are considered non-canonical. In our MIBiG-antiSMASH global analysis, the BGC classes saccharides and “other” were consistently overlooked. Conversely, BGC classes with well-codified domain/protein logics (e.g., NRPS, PKS, and PKS-NRPS hybrids) were successfully detected in most cases, reflecting strong performance in detecting BGCs that align with well-established biosynthetic rules.

This discrepancy also reflected a taxonomy bias: BGCs from *Streptomyces*, a well-known secondary metabolite producer, were more frequently captured by antiSMASH, whereas BGCs from less-studied producers, including diverse Proteobacteria, were often overlooked. GATOR-GC overcomes these limitations by allowing users to define custom protein search parameters, enabling the inclusion of both required and optional proteins. This flexibility bypasses the reliance on pre-defined rules, making it possible to detect non-canonical BGCs that fall outside traditional frameworks. As a result, GATOR-GC complements traditional genome mining tools, expanding the scope of gene cluster discovery, and facilitating the identification of both well-studied and previously uncharacterized gene clusters.

Previous genome mining efforts and manual analysis of gene neighborhoods with ATP-grasp domain-containing protein (MysC) and other MAA-related genes identified 80 MAA BGCs, primarily in Cyanobacteria, with a few exceptions in lil-Proteobacteria and eukaryotes (53). Here using GATOR-GC we expanded the distribution of MAA gene clusters in bacteria, uncovering their widespread presence in Actinomycetota, and highlighting the conservation of genes encoding HAD and TauD/TfdA enzymes. Given their conservation, these enzymes may play a role in the biosynthesis of MAA in Actinomycetota. Unpublished work (63) mentions that the heterologous expression of an MAA operon from *Rhodococcus fascians* D188, containing HAD and TauD enzymes, in *Streptomyces coelicolor M1152* led to the production of shinorine, palythine, and palythine-serine, suggesting that they may function similarly to the nonheme iron-(II)-and 2-oxoglutarate-dependent (Fe/2OG) oxygenase MysH that processes shinorines into palythines (53). TauD may play a role in cleaving the amino acid C-N bond required for palythine biosynthesis. Whether the role of the HAD enzyme, which is highly co-localized with TauD (63), is involved in MAA biosynthesis remains unvalidated. Nonetheless, its consistent presence in actinobacterial MAA gene clusters suggests a potential functional role that warrants further investigation.

Over 70 MAA analogs have already been characterized from taxonomically diverse organisms (51). Using GATOR-GC we were able to computationally identify over 200 MAA gene clusters in this study providing an invaluable resource for the discovery of novel MAA analogs. Given the significant interest in MAAs for the formulation of next-generation sunscreens, our findings not only advance the understanding of MAA biosynthetic diversity but also present promising opportunities for the development of sustainable, bio-based UV-protective compounds.

In addition, targeted genome mining analysis using GATOR-GC expanded the known number of dpd genomic islands from hundreds, as previously reported (55), to tens of thousands. Similar to prior studies, these islands are distributed across phylogenetically diverse bacteria. De Crécy-Lagard et al hypothesized that this broad distribution may be attributed to horizontal gene transfer, supported by their sporadic presence in bacteria, and their insertion at hypervariable loci such as the *leuX* locus in *Salmonella enterica*, a known site for pathogenicity, virulence, and defense genes (64).

These genomic islands, which modify DNA with 7-deazapurines, are thought to act as a self-protective mechanism, shielding DNA from a wide range of restriction enzymes and potentially other defense mechanisms (56). Approximately 27 % of the genomic islands are co-localized with genes involved in the novo biosynthesis of preQ_0_ (e.g., *folE*, *queD*, *queE*, queC), while the remaining islands, lacking these genes, likely either depend on salvaging preQ_0_ from the environment or rely on shared metabolism with other molecules, like queuosine or archaeosine (56). This was demonstrated in the queuosine salvage pathway in *Escherichia coli,* where YhhQ was identified and experimentally validated as a preQ_0_ transporter (65). Notably, YhhQ is present in ∼9 % of the dpd genomic islands, suggesting the potential existence of additional, yet unidentified, transporter families. Further experimental validation is needed to explore this possibility.

GATOR-GC’s ability to detect modular domains in the user’s query proteins proved valuable for targeting gene clusters encoding FK-family compounds. Since an NRPS and a PKS were used as required proteins, GATOR-GC used HMM profiles for their identification, instead of standard DIAMOND-based searches.

GATOR-GC re-discovered gene clusters encoding rapamycin, FK520, FK506, and X1. While these gene clusters share a highly conserved core, including NRPS, PKS, chorismatase, and KCDA, GATOR-GC effectively differentiated their genomic architectures and grouped them according to their chemistry using the GFS. This proximity-scoring system not only captures evolutionary differences by identifying variations across gene clusters but also aids in defining gene cluster boundaries. For example, the only structural difference between FK506 and FK520 is the side chain at C21 (i.e., allyl and ethyl, respectively) (62). If their gene cluster boundaries are considered to lie between the KCDA and O-MT, they are identical in content and organization. However, by incorporating optional proteins encoding the extender units responsible for these side chains, the gene cluster boundaries were extended, allowing GFS to capture the differences. Other more distant FK-family members–such as antascomicins (66), meridamycin (67), and nocardiopsin (68)–were not identified, as their gene clusters lack the gene encoding chorismatase, which was a required search term in our analysis. This reinforces how the inclusion of required and optional genes plays a pivotal role in defining gene cluster identity and grouping related biosynthetic pathways.

In summary, GATOR-GC provides a powerful and flexible framework for exploring gene cluster diversity, overcoming the limitations of other genome mining tools. With customizable search parameters and syntenic analysis, GATOR-GC enhances the discovery of both well-characterized and previously overlooked gene clusters. Its ability to identify gene-level conservation, group similar gene clusters, and reveal evolutionary patterns makes it a valuable tool for bacterial comparative genomics, which is central to uncovering new gene cluster functions and applications.

## Data Availability

The GATOR-GC repository is found at https://github.com/chevrettelab/gator-gc.

## Supplementary Data

## Acknowledgements

We thank Murrel Saldanha and Jordi Weijers for their valuable feedback on the use of GATOR-GC, which helped improve its functionality. We also thank Raquel Dias and Paco Barona-Gómez for their constructive comments and suggestions.

## Author contributions

J.D.D.C.-B, conceptualization, code, design, and development of GATOR-GC, data curation, formal analysis, visualization, writing – original draft, writing – review and editing. A.C, data curation, formal analysis, writing – review, and editing. S.G, data curation, formal analysis, writing – review and editing. Y.D, supervision, writing – review and editing. V.d.C.-L, supervision, writing – review, and editing. M.G.C, conceptualization, code, design and development of GATOR-GC, formal analysis, visualization, funding, supervision, writing – original draft, writing – review and editing.

**Table and Figure Legends**

*Results and Discussion may be combined.

