## Supplementary material for "Targeted genome mining with GATOR-GC maps the evolutionary landscape of biosynthetic diversity": suppl information

**Target Journal: Nucleic Acids Research,** **Computational Biology**

Figures

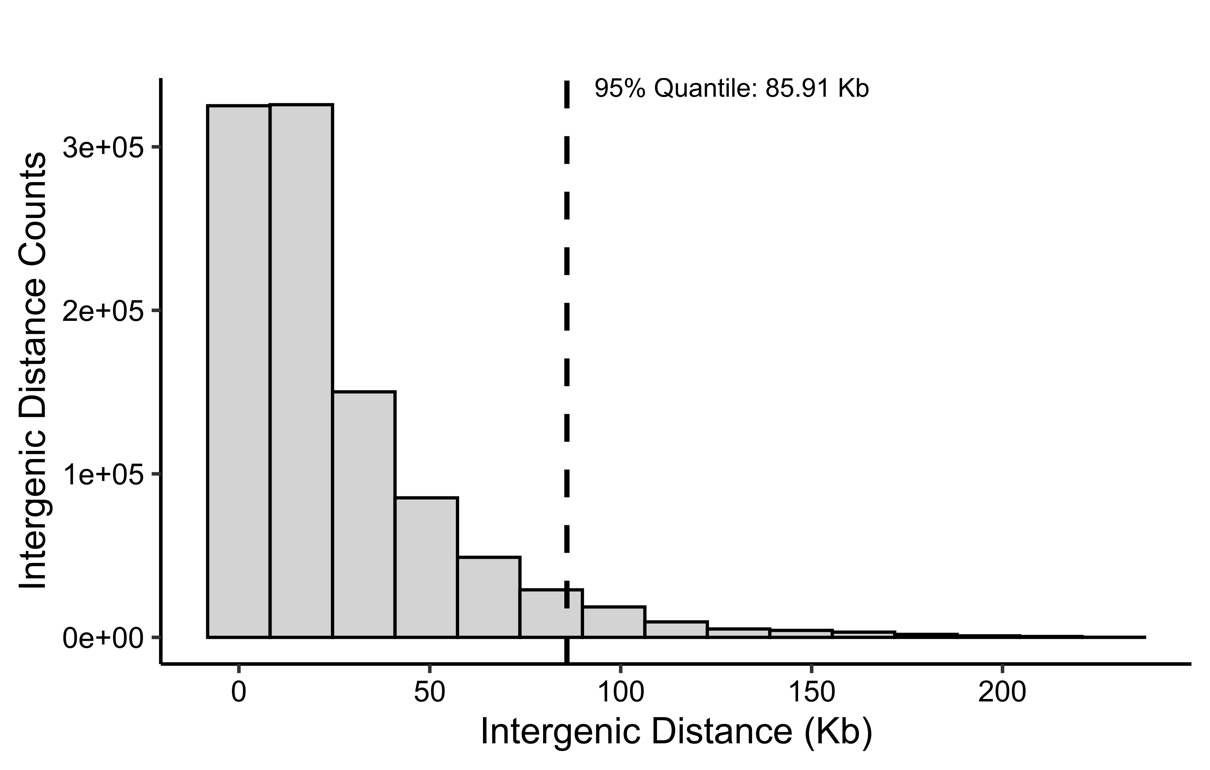

**Supplementary Figure S1. Determination of the required distance parameter value in GATOR-GC.** All intergenic distances between adjacent and non-adjacent genes within MIBiG v3.0 BGCs were analyzed to estimate the optimal distance parameter. The dashed line indicates the 95^th^ percentile of the calculated genomic distances, corresponding to a value of 85.91 Kb. This value was selected as the default parameter for the required distance in GATOR-GC.

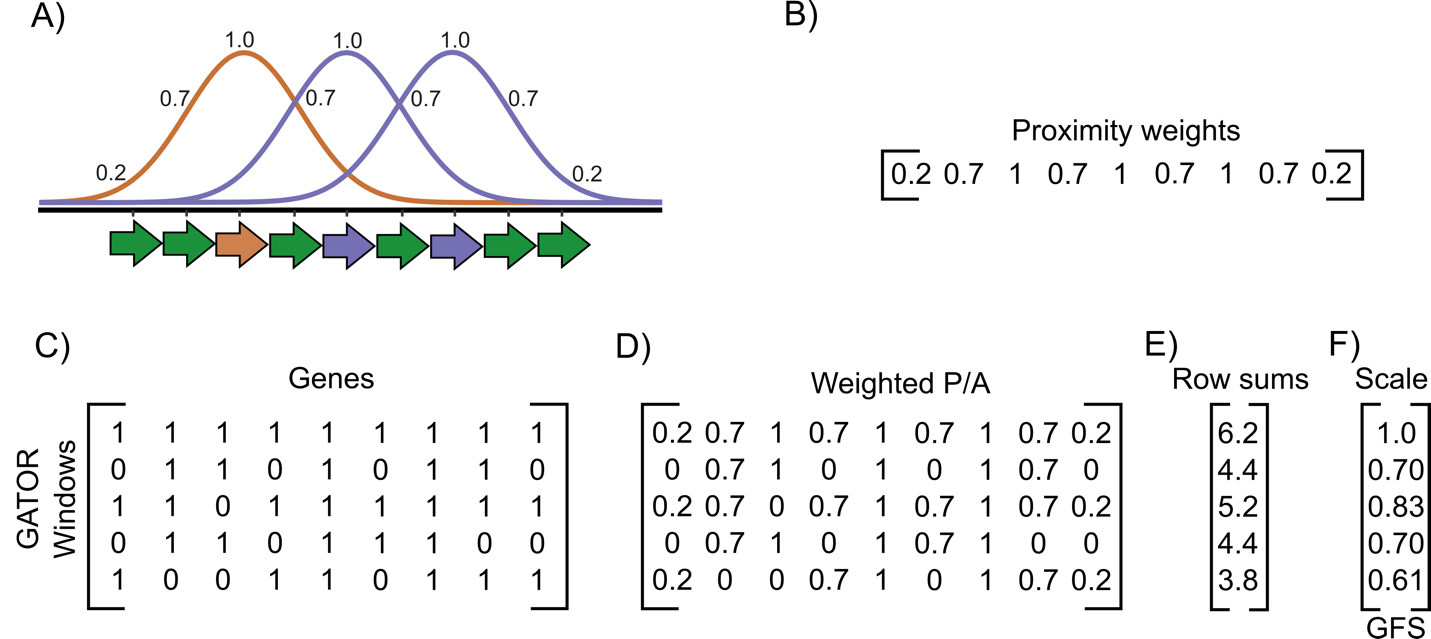

**Supplementary Figure S2. Implementation of the GATOR Focal Scores (GFS).** Given a particular GATOR Focal Window (GFW) that contains two required proteins and one optional protein, three independent weighting distributions are generated (purple for required, orange for optional). These distributions are based on the number of genes within the GFW and the proximity of neighboring genes (A). These proximity weights (B) are then applied to the GATOR windows presence/absence (P/A) tables (C), resulting in weighted P/A tables (D). The weighted values across each GATOR window are summed (E), and these sum values are scaled against the sum value of the GFW to generate the final GFS values (F).

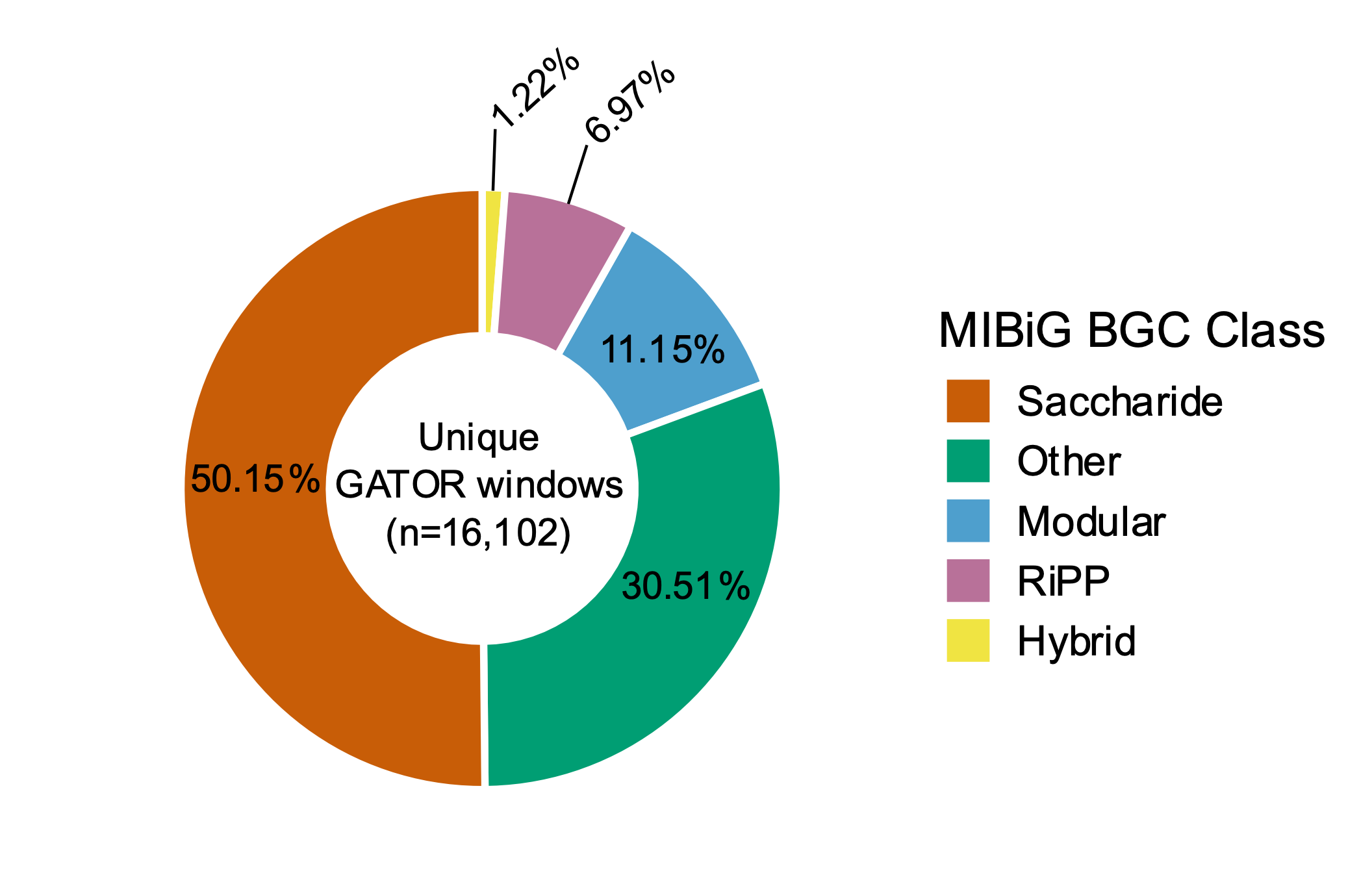

**Supplementary Figure S3. The proportion of deduplicate GATOR windows homologous to the 233 missed bacterial MIBiG v3.0 BGCs.** These windows are classified based on the biosynthetic classes according to the MIBiG. Other include terpenes, alkaloids, and other classes that do not fit into any category. Modular represents NRPS and PKSs.

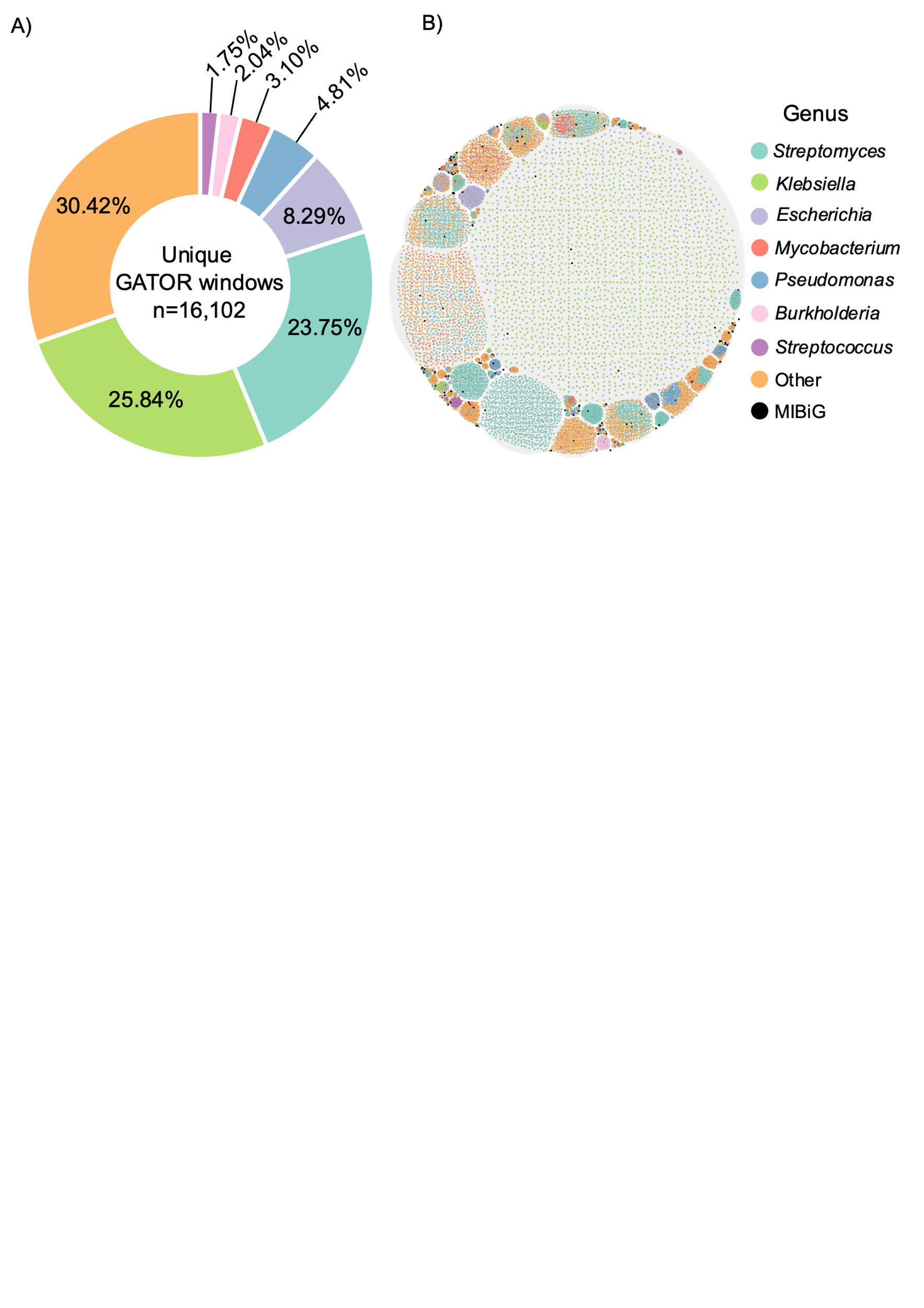
**Supplementary Figure S4. Unique GATOR windows are classified by their genus taxonomy.** The donut plot represents the proportion of the top six genera with the most unique windows (A). GATOR landscape of the BGC families colored by genus taxonomy (B).

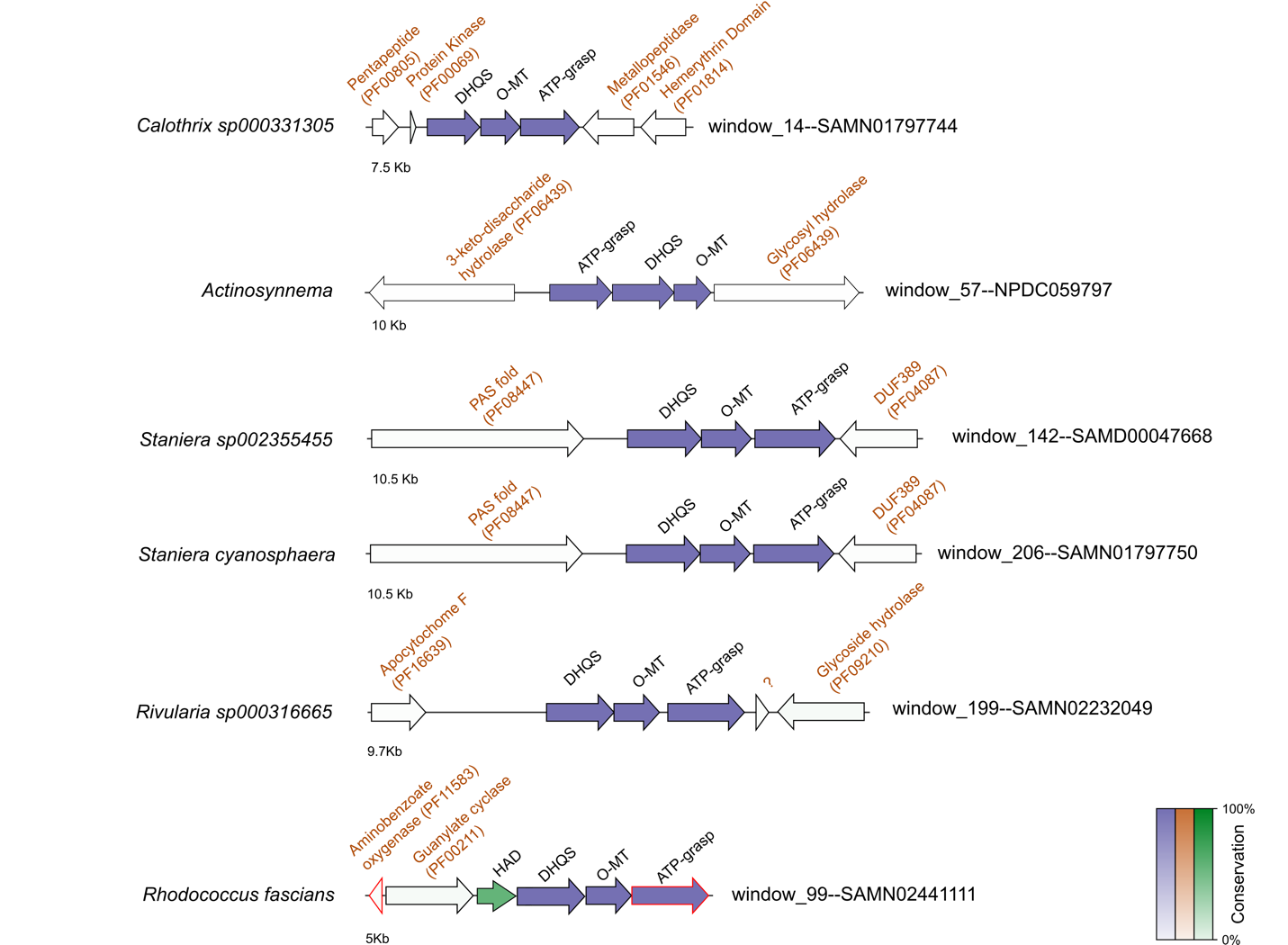

**Supplementary Figure S5. Six GATOR windows in Cyanobacteriota and Actinobacteriota lack similar genes for NRPS-like and d-alanyl-d-alanine ligase enzymes.** PFAM annotations, colored in red, represent genes that are not conserved across the dataset. A question mark (?) indicates protein-encoding genes for which no PFAM domain was detected.

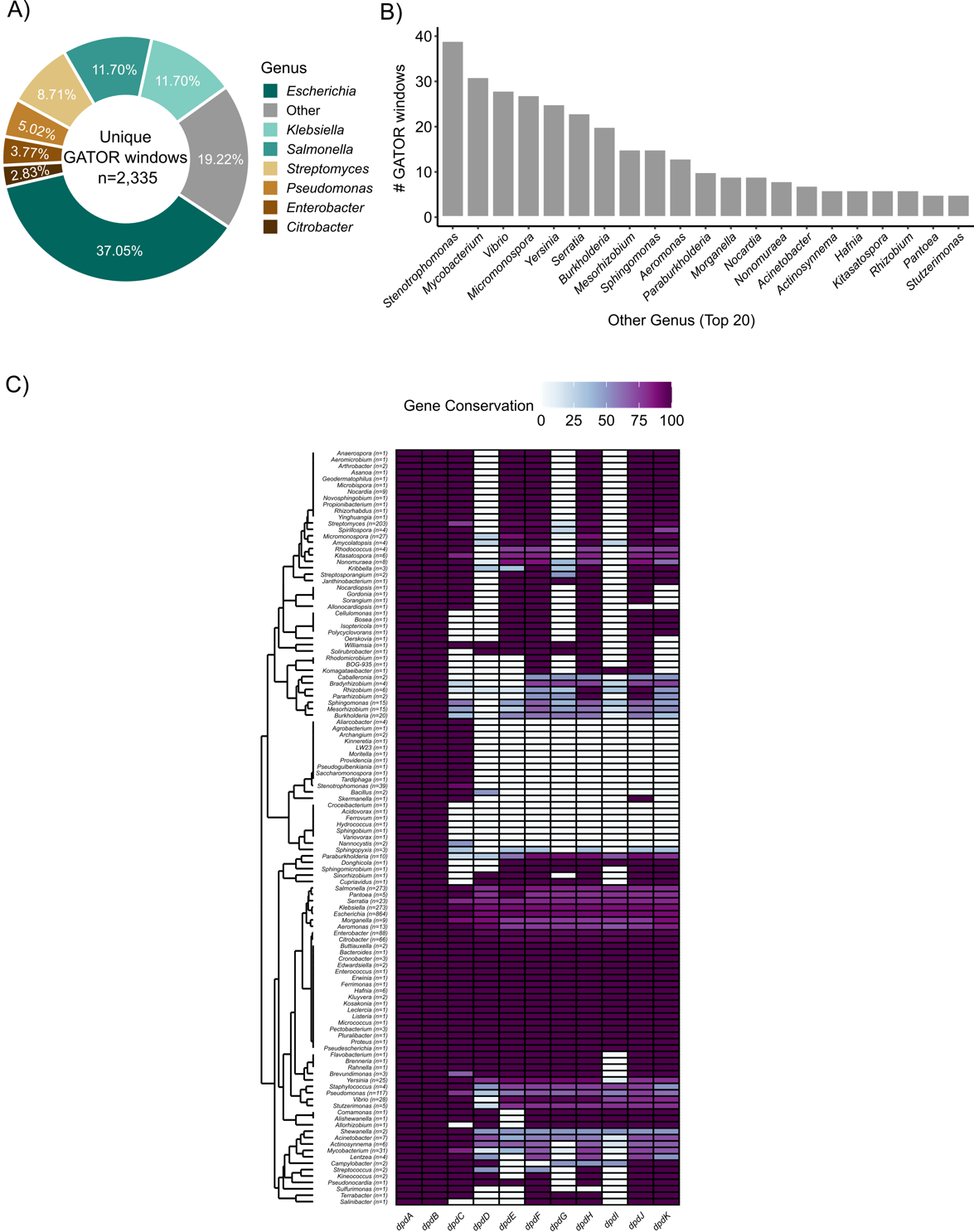

**Supplementary Figure S6. Unique GATOR windows for *dpd* genomic islands, categorized by genus.** The donut plot (A) shows the percentage of windows found in the most frequent genera. Genera with fewer than 40 windows (from the top 20) are shown in (B). The heatmap illustrates the degree of conservation for each gene across all GATOR windows within each genus. Each column in the heatmap represents the genes within the synthesis (*dpdA-C*) and restriction (*dpdD-K*) operons. The dendrogram was constructed using the complete linkage method, based on the presence-absence matrix of the *dpd* genes.

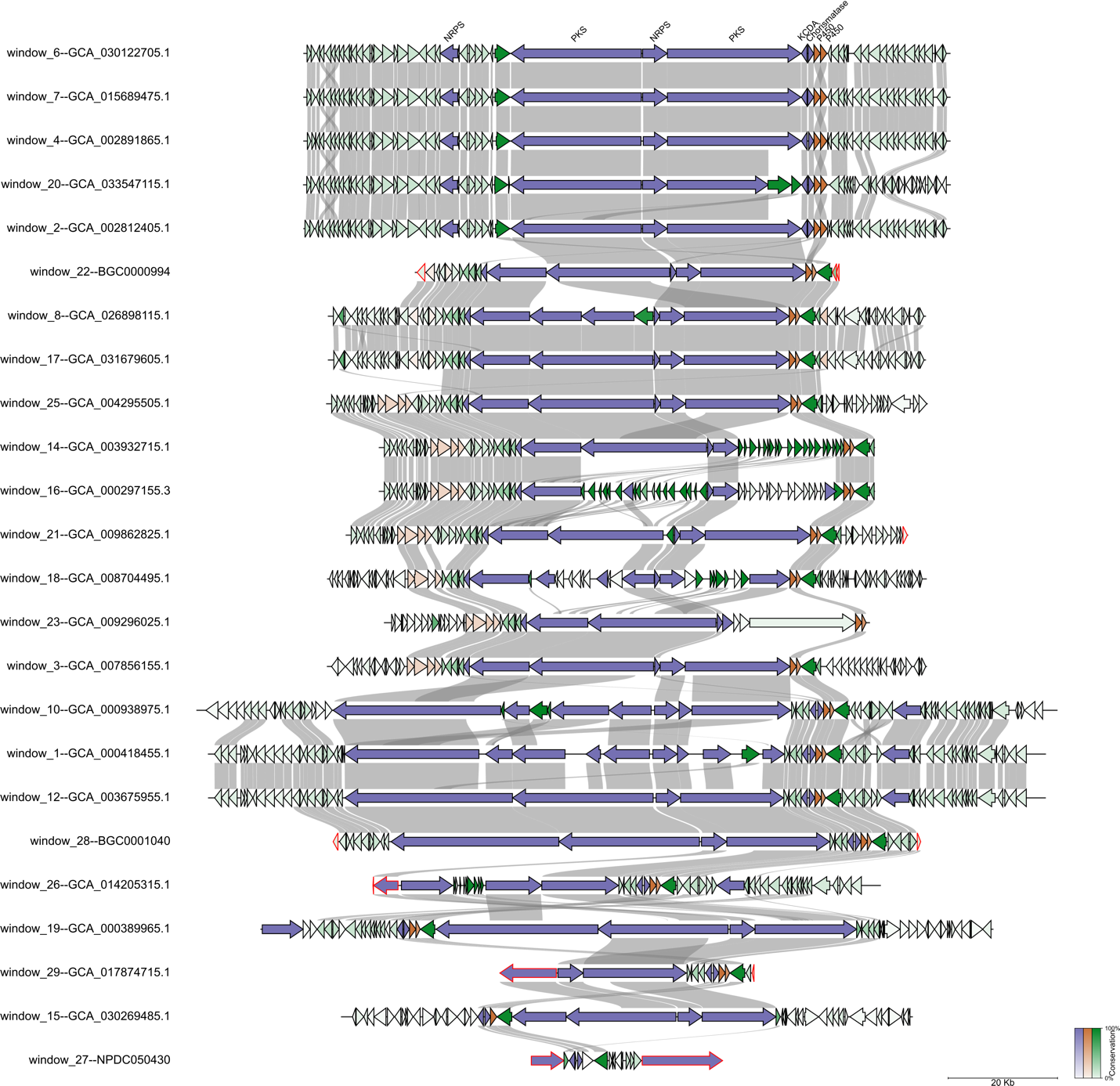

**Supplementary Figure S7. GATOR neighborhoods** **for the unique windows ranked according to the GFS against the X1 BGC.** Required proteins are shown in purple, optional proteins in orange, and other proteins in green. Genes located at contig edges are highlighted with red borders.

Tables

**Supplementary Table S1. Protein search terms for use case analysis.** The table is organized by target gene clusters, with required and optional proteins listed along with their respective protein IDs. The targeted databases for each cluster are also indicated, which include AllTheBacteria (ATB), The Natural Products Discovery Center (NPDC), Actinomycetes from NCBI (ACT), and the Minimum Information about a Biosynthetic Gene Cluster (MIBiG).

| **Target Gene Cluster** | **Required Proteins** | **Protein ID** | **Optional Proteins** | **Protein ID** | **Targeted Database** |
| --- | --- | --- | --- | --- | --- |
| Mycosporine-like amino acids | ATP-Grasp | ABA23461.1 | NRPS-like | ABA23460.1 | ATB, NPDC, MIBiG |
|  | O-MT | ABA23462.1 | D-alanyl-D-alanine ligase | BAY79928.1 |  |
|  | 3-DHQS | ABA23463.1 | SDR dehydrogenase | ANY58988.1 |  |
|  | – | – | Phytanoyl CoA | [BAY79923.1](https://www.ncbi.nlm.nih.gov/protein/1210160838) |  |
| 7-deazapurine DNA modifications | DpdA | WP_001542917.1 | DpdC | WP_223196638.1 | ATB, NPDC, MIBiG |
|  | DpdB | WP_001288809.1 | DpdD | WP_227003010.1 |  |
|  | – | – | DpdE | WP_000431482.1 |  |
|  | – | – | DpdF | WP_001018785.1 |  |
|  | – | – | DpdG | WP_000568475.1 |  |
|  | – | – | DpdH | WP_000209421.1 |  |
|  | – | – | DpdI | WP_000675924.1 |  |
|  | – | – | DpdJ | WP_000580209.1 |  |
|  | – | – | DpdK | WP_000332371.1 |  |
| FK family | PKS | CAA60459.1 | O-MT | CAA60466.1 | ACT, NPDC, MIBiG |
|  | Pipecolate domain | CAA60461.1 | P450 | CAA60469.1 |  |
|  | KCDA | CAA60467.1 | TcsA | ADU56350.1 |  |
|  | Chorismatase | CAA60468.1 | TcsB | ADU56351.1 |  |
|  | – | – | TcsC | ADU56352.1 |  |
|  | – | – | TcsD | ADU56353.1 |  |
|  | – | – | FkbU | [AAF86381.1](https://www.ncbi.nlm.nih.gov/protein/9280383) |  |
|  | – | – | FkbE | [AAF86384.1](https://www.ncbi.nlm.nih.gov/protein/9280386) |  |
|  | – | – | FkbS | [AAF86401.1](https://www.ncbi.nlm.nih.gov/protein/9280403) |  |

**Supplementary Table S2. Assembly accession numbers for representative GATOR windows in dpd genomic islands and FK-family.**

| **Assembly accession numbers** | **Gene cluster** |
| --- | --- |
| SAMN16376653 | 7-deazapurine in DNA (dpd) |
| SAMN07431415 |  |
| SAMN17068888 |  |
| SAMEA104148100 |  |
| SAMN08724136 |  |
| SAMEA4668385 |  |
| SAMN02745261 |  |
| SAMN05890686 |  |
| GCA_030122705.1 | FK-family |
| GCA_031679605.1 |  |
| GCA_004295505.1 |  |
| GCA_007856155.1 |  |
| GCA_003675955.1 |  |
| GCA_000389965.1 |  |
| GCA_030269485.1 |  |

**Supplementary Table S3. Manual curation of the required and optional proteins associated with the missed MIBiG BGCs.**

See the attached file SupplementaryTableS3.xlsx.

**Supplementary Table S4. GATOR-GC results for the overlooked MIBiG BGCs.** This table presents the results of the GATOR-GC analysis for missed MiBIG BGCs, including the corresponding MiBIG BGC type, MiBIG compound, metadata generated by GATOR-GC, and estimates of diversity using the Entropy Shannon. The metadata includes the number of total GATOR windows, the number of unique GATOR Windows (DGW), the mean of the GATOR Focal Scores (GFS), and Entropy values for each taxonomic level (from phylum to species). The table is sorted by the largest number of DGW.

See the attached file SupplementaryTableS4.xlsx.

**Supplementary Table S5**. **GATOR windows detected along with their presence-absence data for enzymes involved in MAA biosynthesis**. For each window, the table shows the window identifier, the genome ID, taxonomy at the species level, and the presence-absence of the associated biosynthetic enzymes.

| **Gator ID** | **Genome ID** | **GTDB Taxonomy** | **DHQS** | **O-MT** | **ATP-grasp** | **D-Ala** | **HAD** | **TauD** | **NRPS-like** | **Phytanoil** | **SDR** |
| --- | --- | --- | --- | --- | --- | --- | --- | --- | --- | --- | --- |
| window_1 | SAMN07376389 | *Rhodococcus sp002259335* | 1 | 1 | 1 | 1 | 1 | 1 | 0 | 0 | 0 |
| window_2 | SAMN03159343 | *Klenkia marina* | 1 | 1 | 1 | 1 | 1 | 1 | 0 | 0 | 0 |
| window_3 | SAMN01797744 | *Calothrix sp000331305* | 1 | 1 | 1 | 0 | 0 | 0 | 0 | 0 | 0 |
| window_4 | SAMN02743366 | *Rhodococcus sp000686025* | 1 | 1 | 1 | 1 | 1 | 1 | 0 | 0 | 0 |
| window_5 | SAMD00153188 | *Mycobacterium madagascariense* | 1 | 1 | 1 | 1 | 1 | 1 | 0 | 0 | 0 |
| window_6 | SAMN07428402 | *Caulobacter vibrioides* | 1 | 1 | 1 | 1 | 1 | 1 | 0 | 0 | 0 |
| window_7 | SAMN02441111 | *Rhodococcus fascians* | 1 | 1 | 1 | 0 | 1 | 0 | 0 | 0 | 0 |
| window_8 | SAMN11484046 | *Nostoc sp013343235* | 1 | 1 | 1 | 1 | 0 | 0 | 0 | 0 | 0 |
| window_9 | SAMN05661030 | *Klenkia taihuensis* | 1 | 1 | 1 | 1 | 1 | 1 | 0 | 0 | 0 |
| window_10 | SAMN14595976 | *Pseudonocardia alni* | 1 | 1 | 1 | 1 | 1 | 1 | 0 | 0 | 0 |
| window_11 | SAMN09240609 | *Rhodococcus kyotonensis_B* | 1 | 1 | 1 | 1 | 1 | 1 | 0 | 0 | 0 |
| window_12 | SAMEA4558873 | *Streptococcus agalactiae* | 1 | 1 | 1 | 1 | 0 | 1 | 0 | 0 | 0 |
| window_13 | SAMD00047668 | *Stanieria sp002355455* | 1 | 1 | 1 | 0 | 0 | 0 | 0 | 0 | 0 |
| window_14 | SAMN10799735 | *Mycobacterium kyogaense* | 1 | 1 | 1 | 1 | 1 | 1 | 0 | 0 | 0 |
| window_15 | SAMN17530832 | *Allobranchiibius huperziae* | 1 | 1 | 1 | 1 | 1 | 1 | 0 | 0 | 0 |
| window_16 | SAMEA3558167 | *Mycobacterium aurum* | 1 | 1 | 1 | 1 | 1 | 1 | 0 | 0 | 0 |
| window_17 | SAMN27922074 | *Nocardioides kribbensis* | 1 | 1 | 1 | 1 | 1 | 1 | 0 | 0 | 0 |
| window_18 | SAMEA113537416 | *Mycobacterium tuberculosis* | 1 | 1 | 1 | 1 | 1 | 1 | 0 | 0 | 0 |
| window_19 | SAMN09468271 | *Brasilonema sp012912105* | 1 | 1 | 1 | 1 | 0 | 0 | 0 | 0 | 0 |
| window_20 | SAMD00242822 | *Trichormus sp019976755* | 1 | 1 | 1 | 0 | 0 | 0 | 1 | 0 | 0 |
| window_21 | SAMN01797750 | *Stanieria cyanosphaera* | 1 | 1 | 1 | 0 | 0 | 0 | 0 | 0 | 0 |
| window_22 | SAMN07376393 | *Rhodococcus fascians_E* | 1 | 1 | 1 | 1 | 1 | 1 | 0 | 0 | 0 |
| window_23 | SAMN05660324 | *Klenkia brasiliensis* | 1 | 1 | 1 | 1 | 1 | 1 | 0 | 0 | 0 |
| window_24 | NPDC080181 | *Rhodococcus fascians* | 1 | 1 | 1 | 1 | 1 | 1 | 0 | 0 | 0 |
| window_25 | SAMN30959949 | *Mycobacterium tuberculosis* | 1 | 1 | 1 | 1 | 1 | 1 | 0 | 0 | 0 |
| window_26 | NPDC078407 | *Rhodococcus* | 1 | 1 | 1 | 1 | 1 | 1 | 0 | 0 | 0 |
| window_27 | SAMN04901515 | *Nostoc sp003326205* | 1 | 1 | 1 | 1 | 0 | 0 | 0 | 1 | 0 |
| window_28 | NPDC026698 | *Pseudonocardia alni* | 1 | 1 | 1 | 1 | 1 | 1 | 0 | 0 | 0 |
| window_29 | SAMD00153213 | *Mycobacterium arabiense* | 1 | 1 | 1 | 1 | 1 | 1 | 0 | 0 | 0 |
| window_30 | SAMN02261353 | *Gloeocapsa sp000332035* | 1 | 1 | 1 | 1 | 0 | 0 | 0 | 0 | 0 |
| window_31 | SAMEA3643012 | *Mycobacterium sp001426545* | 1 | 1 | 1 | 1 | 1 | 1 | 0 | 0 | 0 |
| window_32 | NPDC049306 | *Pseudonocardia alni* | 1 | 1 | 1 | 1 | 1 | 1 | 0 | 0 | 0 |
| window_33 | SAMN10353962 | *Microcystis aeruginosa_E* | 1 | 1 | 1 | 1 | 0 | 0 | 0 | 0 | 0 |
| window_34 | SAMN34233937 | *JAHBTC01 sp002162095* | 1 | 1 | 1 | 1 | 0 | 0 | 0 | 0 | 0 |
| window_35 | NPDC059797 | *Actinosynnema* | 1 | 1 | 1 | 0 | 0 | 0 | 0 | 0 | 0 |
| window_36 | SAMD00573267 | *Mycobacterium* | 1 | 1 | 1 | 1 | 1 | 1 | 0 | 0 | 0 |
| window_37 | SAMN32036409 | *Bradyrhizobium sp003020075* | 1 | 1 | 1 | 1 | 1 | 1 | 0 | 0 | 0 |
| window_38 | SAMN14596006 | *Pseudonocardia alni* | 1 | 1 | 1 | 1 | 1 | 1 | 0 | 0 | 0 |
| window_39 | SAMN14595996 | *Pseudonocardia alni* | 1 | 1 | 1 | 1 | 1 | 1 | 0 | 0 | 0 |
| window_40 | SAMN02569990 | *Rhodococcus sp002259335* | 1 | 1 | 1 | 1 | 1 | 1 | 0 | 0 | 0 |
| window_41 | SAMD00197826 | *Chroococcus sp017312365* | 1 | 1 | 1 | 1 | 0 | 0 | 0 | 1 | 0 |
| window_42 | SAMN13734961 | *Rhodococcus kyotonensis_B* | 1 | 1 | 1 | 1 | 1 | 1 | 0 | 0 | 0 |
| window_43 | SAMN02441755 | *Pelatocladus sp000447295* | 1 | 1 | 1 | 0 | 0 | 0 | 1 | 0 | 0 |
| window_44 | SAMN05660199 | *Klenkia soli* | 1 | 1 | 1 | 1 | 1 | 1 | 0 | 0 | 0 |
| window_45 | SAMN05216377 | *Pseudonocardia oroxyli* | 1 | 1 | 1 | 1 | 1 | 1 | 0 | 0 | 0 |
| window_46 | SAMN04901525 | *Dendronalium minutum* | 1 | 1 | 1 | 1 | 0 | 0 | 0 | 0 | 0 |
| window_47 | SAMN03165654 | *Trichormus* | 1 | 1 | 1 | 1 | 0 | 0 | 0 | 0 | 0 |
| window_48 | SAMN13620235 | *Komarekiella sp010091925* | 1 | 1 | 1 | 1 | 0 | 0 | 0 | 0 | 0 |
| window_49 | NPDC000705 | *Actinosynnema mirum* | 1 | 1 | 1 | 1 | 0 | 0 | 0 | 0 | 0 |
| window_50 | SAMN11588057 | *Trichormus sp013393945* | 1 | 1 | 1 | 0 | 0 | 0 | 1 | 0 | 0 |
| window_51 | SAMN34233953 | *Blastomonas fulva* | 1 | 1 | 1 | 1 | 0 | 0 | 0 | 0 | 0 |
| window_52 | SAMD00153170 | *Mycobacterium gallinarum* | 1 | 1 | 1 | 1 | 1 | 1 | 0 | 0 | 0 |
| window_53 | SAMN10353975 | *Microcystis sp002282935* | 1 | 1 | 1 | 1 | 0 | 0 | 0 | 0 | 0 |
| window_54 | SAMEA3642918 | *Nocardioides kribbensis* | 1 | 1 | 1 | 1 | 1 | 1 | 0 | 0 | 0 |
| window_55 | SAMN07658331 | *Mycobacterium* | 1 | 1 | 1 | 1 | 1 | 1 | 0 | 0 | 0 |
| window_56 | SAMN02194776 | *Rhodococcus fascians_E* | 1 | 1 | 1 | 1 | 1 | 1 | 0 | 0 | 0 |
| window_57 | SAMD00156906 | *Sphaerospermopsis kisseleviana_A* | 1 | 1 | 1 | 1 | 0 | 0 | 0 | 0 | 0 |
| window_58 | BGC0001615 | MiBIG | 1 | 1 | 1 | 0 | 0 | 0 | 1 | 0 | 1 |
| window_59 | SAMEA113538554 | *Mycobacterium tuberculosis* | 1 | 1 | 1 | 1 | 1 | 1 | 0 | 0 | 0 |
| window_60 | SAMN07658303 | *Mycobacterium tokaiense* | 1 | 1 | 1 | 1 | 1 | 1 | 0 | 0 | 0 |
| window_61 | SAMEA3642943 | *Klenkia sp001424485* | 1 | 1 | 1 | 1 | 1 | 1 | 0 | 0 | 0 |
| window_62 | SAMN09074697 | *Williamsia limnetica* | 1 | 1 | 1 | 1 | 1 | 1 | 0 | 0 | 0 |
| window_63 | SAMN10353977 | *CAILKP01 sp903834785* | 1 | 1 | 1 | 1 | 0 | 0 | 0 | 0 | 0 |
| window_64 | SAMN05921184 | *Microcystis panniformis* | 1 | 1 | 1 | 1 | 0 | 0 | 0 | 0 | 0 |
| window_65 | SAMN06311396 | *Nodularia sp015207755* | 1 | 1 | 1 | 1 | 0 | 0 | 0 | 0 | 0 |
| window_66 | NPDC079359 | *Rhodococcus sp002259485* | 1 | 1 | 1 | 1 | 1 | 1 | 0 | 0 | 0 |
| window_67 | SAMN19550771 | *Mycobacterium kyogaense* | 1 | 1 | 1 | 1 | 1 | 1 | 0 | 0 | 0 |
| window_68 | SAMEA3643095 | *Aeromicrobium sp001426485* | 1 | 1 | 1 | 1 | 1 | 0 | 0 | 0 | 0 |
| window_69 | BGC0000427 | MiBIG | 1 | 1 | 1 | 0 | 0 | 0 | 1 | 0 | 0 |
| window_70 | SAMN02261327 | *Calothrix sp000317435* | 1 | 1 | 1 | 1 | 0 | 0 | 0 | 0 | 0 |
| window_71 | SAMN02261335 | *Microcoleus sp000317475* | 1 | 1 | 1 | 1 | 0 | 0 | 0 | 0 | 0 |
| window_72 | NPDC047994 | *Pseudonocardia alni* | 1 | 1 | 1 | 1 | 1 | 1 | 0 | 0 | 0 |
| window_73 | SAMN02261253 | *Oscillatoria sp000332335* | 1 | 1 | 1 | 0 | 0 | 0 | 1 | 0 | 0 |
| window_74 | SAMN11587828 | *Nostoc sp013393905* | 1 | 1 | 1 | 1 | 0 | 0 | 0 | 0 | 0 |
| window_75 | SAMEA3642966 | *Marmoricola sp001424755* | 1 | 1 | 1 | 1 | 1 | 1 | 0 | 0 | 0 |
| window_76 | SAMN02261257 | *Pelatocladus prolificus* | 1 | 1 | 1 | 0 | 0 | 0 | 1 | 0 | 0 |
| window_77 | NPDC060226 | *Streptomyces* | 1 | 1 | 1 | 1 | 1 | 1 | 0 | 0 | 0 |
| window_78 | SAMN14846999 | *Mycobacterium hippocampi_A* | 1 | 1 | 1 | 1 | 1 | 1 | 0 | 0 | 0 |
| window_79 | SAMN01797701 | *Trichormus sp000316645* | 1 | 1 | 1 | 0 | 0 | 0 | 1 | 0 | 0 |
| window_80 | SAMN13190112 | *Mycobacterium* | 1 | 1 | 1 | 1 | 1 | 1 | 0 | 0 | 0 |
| window_81 | SAMN02256421 | *Actinomycetospora chiangmaiensis* | 1 | 1 | 1 | 1 | 1 | 0 | 0 | 0 | 0 |
| window_82 | SAMN17172209 | *Trichormus variabilis* | 1 | 1 | 1 | 0 | 0 | 0 | 1 | 0 | 0 |
| window_83 | SAMN05445060 | *Williamsia_B sterculiae* | 1 | 1 | 1 | 1 | 1 | 1 | 0 | 0 | 0 |
| window_84 | SAMN04453661 | *Nostoc sp001712795* | 1 | 1 | 1 | 1 | 0 | 0 | 0 | 0 | 0 |
| window_85 | SAMN14595995 | *Pseudonocardia alni* | 1 | 1 | 1 | 1 | 0 | 1 | 0 | 0 | 0 |
| window_86 | NPDC050041 | *Mycobacterium* | 1 | 1 | 1 | 1 | 1 | 1 | 0 | 0 | 0 |
| window_87 | SAMD00153209 | *Mycobacterium sediminis* | 1 | 1 | 1 | 1 | 1 | 1 | 0 | 0 | 0 |
| window_88 | NPDC057246 | *Rhodococcus kroppenstedtii* | 1 | 1 | 1 | 1 | 1 | 1 | 0 | 0 | 0 |
| window_89 | SAMN05967630 | *Nodularia spumigena* | 1 | 1 | 1 | 1 | 0 | 0 | 0 | 0 | 0 |
| window_90 | SAMN02569995 | *Rhodococcus sp002259405* | 1 | 1 | 1 | 1 | 1 | 1 | 0 | 0 | 0 |
| window_91 | SAMN34233941 | *Erythrobacter sp019750455* | 1 | 1 | 1 | 1 | 0 | 0 | 0 | 0 | 0 |
| window_92 | NPDC002237 | *Pseudonocardia alni* | 1 | 1 | 1 | 1 | 1 | 1 | 0 | 0 | 0 |
| window_93 | SAMN17530815 | *Allobranchiibius sp017565405* | 1 | 1 | 1 | 1 | 1 | 1 | 0 | 0 | 0 |
| window_94 | SAMEA7528687 | *Mycobacterium tuberculosis* | 1 | 1 | 1 | 1 | 1 | 1 | 0 | 0 | 0 |
| window_95 | SAMN13734960 | *Rhodococcus fascians* | 1 | 1 | 1 | 1 | 1 | 1 | 0 | 0 | 0 |
| window_96 | SAMN34233943 | *Dolichospermum sp000312705* | 1 | 1 | 1 | 1 | 0 | 0 | 0 | 0 | 0 |
| window_97 | SAMN05660695 | *Williamsia_A maris* | 1 | 1 | 1 | 1 | 1 | 1 | 0 | 0 | 0 |
| window_98 | SAMEA3642894 | *Rhodococcus sp001426185* | 1 | 1 | 1 | 1 | 1 | 1 | 0 | 0 | 0 |
| window_99 | SAMN20254267 | *Mycobacterium rufum* | 1 | 1 | 1 | 1 | 1 | 1 | 0 | 0 | 0 |
| window_100 | SAMN01797693 | *Cylindrospermum stagnale* | 1 | 1 | 1 | 1 | 0 | 0 | 0 | 0 | 0 |
| window_101 | SAMN02261341 | *Anabaena sp000332135* | 1 | 1 | 1 | 1 | 0 | 0 | 0 | 0 | 0 |
| window_102 | SAMEA3642941 | *Klenkia sp001424455* | 1 | 1 | 1 | 1 | 1 | 1 | 0 | 0 | 0 |
| window_103 | SAMN02569994 | *Priestia zanthoxyli* | 1 | 1 | 1 | 1 | 1 | 1 | 0 | 0 | 0 |
| window_104 | SAMN05661029 | *Williamsia_A deligens* | 1 | 1 | 1 | 1 | 1 | 1 | 0 | 0 | 0 |
| window_105 | SAMN18296527 | *Crocosphaera watsonii* | 1 | 1 | 1 | 1 | 0 | 0 | 0 | 0 | 0 |
| window_106 | SAMN15375537 | *Nodularia spumigena* | 1 | 1 | 1 | 1 | 0 | 0 | 0 | 0 | 0 |
| window_107 | SAMN07604078 | *Trichormus sp002896875* | 1 | 1 | 1 | 0 | 0 | 0 | 1 | 0 | 0 |
| window_108 | SAMEA1530731 | *Mycobacterium tuberculosis* | 1 | 1 | 1 | 1 | 1 | 1 | 0 | 0 | 0 |
| window_109 | SAMN02569997 | *Rhodococcus fascians* | 1 | 1 | 1 | 1 | 1 | 1 | 0 | 0 | 0 |
| window_110 | SAMN13242330 | *Nostoc muscorum_A* | 1 | 1 | 1 | 1 | 0 | 0 | 0 | 0 | 0 |
| window_111 | SAMN05928532 | *Microcystis panniformis_A* | 1 | 1 | 1 | 1 | 0 | 0 | 0 | 0 | 0 |
| window_112 | SAMN02232049 | *Rivularia sp000316665* | 1 | 1 | 1 | 0 | 0 | 0 | 0 | 0 | 0 |
| window_113 | SAMN14595967 | *Pseudonocardia alni* | 1 | 1 | 1 | 1 | 1 | 1 | 0 | 0 | 0 |
| window_114 | SAMN02569999 | *Rhodococcus fascians* | 1 | 1 | 1 | 1 | 1 | 1 | 0 | 0 | 0 |
| window_115 | SAMN07660175 | *Mycobacterium iranicum* | 1 | 1 | 1 | 1 | 1 | 1 | 0 | 0 | 0 |
| window_116 | NPDC076796 | *Rhodococcus sp000813105* | 1 | 1 | 1 | 1 | 1 | 1 | 0 | 0 | 0 |
| window_117 | SAMN12025127 | *Mycobacterium iranicum_B* | 1 | 1 | 1 | 1 | 1 | 1 | 0 | 0 | 0 |
| window_118 | SAMN02744079 | *Hyella* | 1 | 1 | 1 | 1 | 0 | 0 | 0 | 0 | 0 |
| window_119 | SAMD00153184 | *Mycobacterium psychrotolerans* | 1 | 1 | 1 | 1 | 1 | 1 | 0 | 0 | 0 |
| window_120 | SAMN15196599 | *Nostoc edaphicum_A* | 1 | 1 | 1 | 1 | 0 | 0 | 0 | 0 | 0 |
| window_121 | NPDC000082 | *Actinosynnema pretiosum* | 1 | 1 | 1 | 1 | 0 | 0 | 0 | 0 | 0 |
| window_122 | SAMN08778927 | *Rhodococcus* | 1 | 1 | 1 | 1 | 1 | 1 | 0 | 0 | 0 |
| window_123 | SAMN05443575 | *Jatrophihabitans endophyticus* | 1 | 1 | 1 | 1 | 1 | 1 | 0 | 0 | 0 |
| window_124 | SAMEA3642936 | *Williamsia_A herbipolensis* | 1 | 1 | 1 | 1 | 1 | 1 | 0 | 0 | 0 |
| window_125 | SAMN11658418 | *Mycobacterium* | 1 | 1 | 1 | 1 | 1 | 1 | 0 | 0 | 0 |
| window_126 | NPDC048788 | *Pseudonocardia alni* | 1 | 1 | 1 | 1 | 1 | 1 | 0 | 0 | 0 |
| window_127 | SAMD00042755 | *Trichormus sp001548375* | 1 | 1 | 1 | 0 | 0 | 0 | 1 | 0 | 0 |
| window_128 | SAMD00545464 | *Mycobacterium* | 1 | 1 | 1 | 1 | 1 | 1 | 0 | 0 | 0 |
| window_129 | NPDC023957 | *Pseudonocardia alni* | 1 | 1 | 1 | 1 | 1 | 1 | 0 | 0 | 0 |
| window_130 | NPDC057297 | *Rhodococcus sp002259415* | 1 | 1 | 1 | 1 | 1 | 1 | 0 | 0 | 0 |
| window_131 | SAMEA1035219 | *Rhodococcus sp002259485* | 1 | 1 | 1 | 1 | 1 | 1 | 0 | 0 | 0 |

**Supplementary Table S6. Unique dpd genomic islands with their taxonomic information and gene-level presence-absence of *dpd* genes.**

See the attached file SupplementaryTableS6.xlsx.

**Supplementary Table S7. Variability of 133 *dpd* gene architectures in bacterial genomic Islands.** This table details the different architectures of *dpd* genes identified within the deduplicated set of 2,335 genomic islands. Each unique architecture is listed along with the number of genomic islands in which it is found, and the corresponding percentage of the total genomic islands.

See the attached file SupplementaryTableS7.xlsx.

**Supplementary Table S8. Prevalence of *dpd* genes in dpd genomic islands**. This table summarizes the presence of *dpd* genes across 2,335 deduplicated genomic islands. Each gene is listed along with its corresponding protein ID, the number of genomic islands in which it is present, and the percentage of genomic islands containing that single gene. Additionally, the number of genes with more than one copy per genomic island is shown, along with the total number of additional copies across the dataset.

| ***dpds*** | **Single occurrence** | **Percentage** | **Multiple occurrences** | **Total of additional copies** |
| --- | --- | --- | --- | --- |
| dpdA\|WP_001542917.1 | 2335 | 100 | 2369 | 34 |
| dpdB\|WP_001288809.1 | 2335 | 100 | 3136 | 801 |
| dpdC\|WP_223196638.1 | 2151 | 92.12 | 2158 | 7 |
| dpdF\|WP_000431482.1 | 2015 | 86.3 | 2054 | 39 |
| dpdJ\|WP_000675924.1 | 2006 | 85.91 | 2035 | 29 |
| dpdH\|WP_000568475.1 | 2004 | 85.82 | 2023 | 19 |
| dpdE\|WP_227003010.1 | 1983 | 84.93 | 2007 | 24 |
| dpdK\|WP_000580209.1 | 1942 | 83.17 | 1943 | 1 |
| dpdG\|WP_001018785.1 | 1714 | 73.4 | 1722 | 8 |
| dpdI\|WP_000209421.1 | 1580 | 67.67 | 1583 | 3 |
| dpdD\|WP_000332371.1 | 1572 | 67.32 | 1586 | 14 |

**Supplementary Table S9. Prevalence of PFAM domains in dpd genomic islands**. This table summarizes the presence of unique PFAM domains across 2,335 unique genomic islands. Each domain is listed along with its corresponding PFAM annotation, the number of genomic islands in which it is present, and the percentage of genomic islands containing that single domain. Domains highlighted in yellow represent those that cluster with *dpd* genes in over 20% of the islands.

See the attached file SupplementaryTableS9.xlsx.
